# Higher stomatal density and smaller stomatal size associate with higher grain yield in winter wheat

**DOI:** 10.64898/2026.11.04.764249

**Authors:** Elena Ivandi, Pirko Jalakas, Ingmar Tulva, Mati Koppel, Kajal Samantara, Banafsheh Khalegh Doust, Anne Ingver, Merlin Haljak, Reine Koppel, Ebe Merilo, Hanna Hõrak

## Abstract

In wheat (*Triticum aestivum L*.), flag leaf is crucial in determining grain-filling rate and final grain yield. Stomata, microscopic pores on plant surfaces, balance CO_2_ uptake and evaporative water loss; their numbers and aperture sizes depend on environmental factors. Previous studies have reported associations between stomatal traits and grain yield; mostly under controlled conditions. Our study aimed to investigate flag leaf stomatal anatomical and physiological traits, and their relationships with winter wheat grain yield under field conditions. We measured stomatal density and size, and stomatal conductance on adaxial and abaxial flag leaf surfaces in a three-year field study with twenty-five winter wheat varieties. Stomatal density ratio remained constant despite different stomatal densities occurring in different years, indicating coordinated adjustments of stomatal numbers between leaf surfaces in response to environmental conditions. Stomatal conductance ratios varied widely, indicating leaf-surface specific stomatal responses to environmental conditions. Grain yield was positively associated with stomatal density and negatively with stomatal size. Flag leaf removal reduced grain weight per spike and did not affect grain numbers per spike, unlike in spring wheat. Our field study provides valuable insights into the relationships between winter wheat stomatal traits and grain yield that can be used in breeding.

**Highlight:** We found coordinated adjustment of wheat stomatal densities, but independent stomatal responses to environmental factors on the adaxial and abaxial leaf surfaces, whereas stomatal density positively associated with grain yield.

## 1. Introduction

Wheat (*Triticum aestivum L*.) is one of the most widely grown crops and plays a crucial role in global food security (Erenstein *et al*., 2022). In 2025, the global wheat production was 843.8 Mt (‘Wheat | USDA Foreign Agricultural Service’, 2026). The flag leaf is the major organ in wheat canopy, determining grain-filling rate, yield components such as grain numbers per spike and grain weight per spike, and overall grain yield by supplying photoassimilates to developing grains after anthesis (Melahat Avci, 2005; Ali *et al*., 2010; Racz *et al*., 2022). As the topmost youngest leaf, the flag leaf absorbs more light compared with lower, shaded leaves. Removal of the flag leaf negatively affects grain yield in spring and winter wheat and forces lower leaves to compensate and to supply photoassimilates to grains (Patterson and Ohm, 1975; Mahmood *et al*., 1991; Melahat Avci, 2005; Racz *et al*., 2022). Since most of the previous studies have been conducted in warm climates, the importance of the flag leaf for the formation of grain yield components under cooler conditions of Northern Europe remains unclear.

Glaucousness, a characteristic associated with adaptation to water-limited and heat-stressed conditions, manifests as a visible waxy epicuticular layer with a dull white to bluish color on leaves and spikes (Johnson *et al*., 1983; Merah *et al*., 2000; Guo *et al*., 2016; Bi *et al*., 2017). Epicuticular waxes of glaucous cultivars serve as a barrier preventing non-stomatal water loss, leading to higher water use efficiency, increased leaf longevity, and lower leaf temperature, potentially due to increased reflectance (Johnson *et al*., 1983; Richards *et al*., 1986). Glaucous cultivars can have higher yields both in drought conditions (Merah *et al*., 2000) and when water is not limiting (Johnson *et al*., 1983).

Stomata are microscopic pores on plant surfaces (leaves, stems) that balance CO_2_ uptake and evaporative water loss, exerting significant control on the world’s water and carbon cycles (Hetherington and Woodward, 2003). Depending on species, stomata can be present on one leaf surface or be distributed between both leaf surfaces. Most plants are hypostomatous, meaning that they have stomata only on the lower (abaxial) leaf surface (Muir, 2015). In amphistomatous plants, stomata are formed on both leaf surfaces, whereas they develop and function at least partly independently (Hõrak, 2026). Stomatal density (SD) in most amphistomatous species tends to be higher on the abaxial (lower) compared with the adaxial (upper) surface (Pemadasa, 1979). The exception is the *Poaceae* family: their SD is often equal on both leaf surfaces, or higher on the adaxial surface, as in wheat (Pemadasa, 1979; Wall *et al*., 2022; Samantara *et al*., 2025), where adaxial surface also contributes more to leaf gas-exchange (Wall *et al*., 2022; Chaplin *et al*., 2026a; Chaplin *et al*., 2026b). Adaxial and abaxial stomata in amphistomatous plants respond independently to environmental stimuli, such as light intensity (Wang *et al*., 2008; Wall *et al*., 2022; Wei *et al*., 2025). Higher adaxial to abaxial stomatal density ratio (SR) has been associated with faster stomatal response kinetics (Woning et al., 2026; Yang et al., 2026). Having more stomata on the adaxial leaf surface increases photosynthetic efficiency and CO_2_ uptake (Xiong and Flexas, 2020), but it also may increase evaporative water loss and susceptibility to stomata-entering pathogens (e.g., rust fungi) (McKown *et al*., 2014; Muir, 2015; Drake *et al*., 2019). Yield losses caused by pathogens are associated with lodging, decreased nutrient availability, and lower thousand grain weight, and numbers of grain per spike (Figueroa *et al*., 2018; Khanfri *et al*., 2018). Higher SDs may lead to increased susceptibility to stomata-entering pathogens (Dutton *et al*., 2019; Tateda *et al*., 2019). Thus, leaf surface-specific stomatal traits may influence CO_2_ uptake, water loss, and pathogen entry, ultimately affecting yield under field conditions.

Wheat stomatal traits are associated with grain yield, but specific relationships may depend on growth stage and environment. For example, a positive correlation was found between abaxial SD and grain yield (Yang *et al*., 2022; Du *et al*., 2026), or both adaxial and abaxial SD and grain yield (Yousufzai *et al*., 2009) in wheat. Other studies have reported a negative relationship between adaxial SD and grain yield (Samantara *et al*., 2025), or found no correlation between stomatal traits and grain yield (Ledent and Jouret, 1978; Shahinnia *et al*., 2016; Chaplin *et al*., 2026a). Thus, stomatal traits may have breeding potential in wheat but further characterisation of leaf surface-specific stomatal anatomy and physiology under field conditions is needed to better understand the variation in wheat stomatal traits and their contribution to yield determination.

This study aimed to (i) investigate winter wheat flag leaf stomatal anatomical and physiological traits under field conditions over three years, (ii) examine the relationships between flag leaf adaxial and abaxial stomatal traits and grain yield of winter wheat, and (iii) determine the effect of flag leaf removal on yield components in winter wheat and spring wheat.

## 2. Material and methods

### 2.1. Plant material and field experiments

Field trials were conducted at the Centre of Estonian Rural Research and Knowledge (METK, Estonia), Jõgeva, Estonia (58°45’N, 26°24’E) in 2023-2025 seasons. Twenty-five winter wheat varieties (Table S1) in three replications were sown (plot size of 9 m^2^, and row spacings of 12.5 cm) on 9 September 2022; 11 September 2023, and on 9 September 2024. Fungicide treatment was not applied. Half of the varieties included in the trial were early-maturing and the other half were late-maturing (Table S1). In 2025, we included ten spring wheat varieties with three replications (three plots per genotype) in the flag leaf removal experiment as a comparison to winter wheat. Plot size was the same as for winter wheat. Spring wheat was sown on 28 April 2025. Field trials were carried out following a randomized complete block design.

### 2.2. Meteorological data

In 2023–2024, May-June were relatively dry, followed by rainy July (Fig. 1). The wettest year was 2025, when precipitation was above long-period average (LPA, 1991-2020) for most of the spring and summer months. In 2023–2024, temperatures were high from May to August, while in 2025, temperatures were below LPA during the major growth period in May and June, and exceeded the LPA in the main grain filling period in July (Fig. 1).

**Figure 1.**
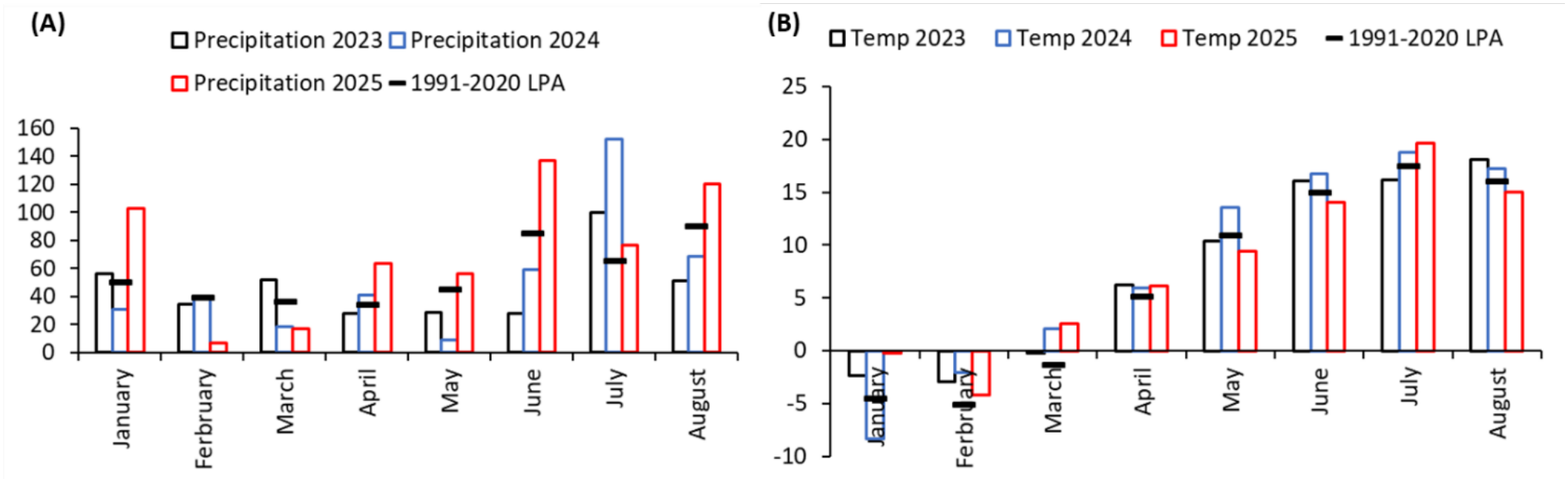
Weather data in Jõgeva in 2023–2025. (A) Total precipitation from January to August in 2023–2025 in Jõgeva in comparison with the respective long-period average (LPA, 1991– 2020); (B) Temperatures in 2023–2025 in Jõgeva in comparison with respective long-period average (LPA, 1991–2020). Weather data from Estonian Environment Agency (Station location in Jõgeva N 58°44′59″, E 26°24′54″) Alt text: Graphs labelled A–B, A is showing precipitation in Jõgeva in 2023–2025 from January to August, and B is showing temperatures in Jõgeva in 2023–2025 from January to August.

### 2.3. Flag leaf traits

Since half of the varieties included in the study were early-maturing and the other half were late-maturing (Table S1), the flag leaf measurements were taken twice, about one week apart (9 and 16 June in 2023; 7 and 11 June in 2024; 16 and 26 June in 2025; flag leaves of early varieties were measured and detached for anatomical measurements on the first and those of late varieties on the second date each year) at the end of the heading stage (BBCH 59), when flag leaves had reached full size. Six plants from each plot were randomly selected for flag leaf stomatal measurements and flag leaf removal, and an additional six plants with intact flag leaves were chosen as control plants for yield and yield component determination at the final harvest. The abaxial and adaxial stomatal conductance (g_s_) of the central part of flag leaves (avoiding the midrib) was measured using a LI-600 (LI-COR) hand-held porometer. In further data analysis, total g_s_ was calculated as the sum of abaxial and adaxial g_s_ and g_s_ ratio as the ratio of adaxial g_s_ to abaxial g_s_. After g_s_ determination, measured flag leaves were removed and taken to the laboratory for flag leaf area and stomatal anatomical characteristics measurements. In the laboratory, the flag leaves were photographed, and flag leaf areas (FLA, cm^2^) and widths were measured using ImageJ software (Schneider *et al*., 2012). At late wax maturity (BBCH 89), before mechanical harvesting, 12 marked plants (six with flag leaf removed and six control plants) were harvested manually from each plot for determination of plant yield components of the main shoot (grain number per spike and grain weight per spike). After that, winter wheat was harvested mechanically between 28 July and 8 August, 2023, between 21 July and 2 August in 2024, and between 31 July and 8 August in 2025. After mechanical harvest, the thousand grain weight (TGW, g) of winter wheat was measured, grain yield (kg ha^-1^) was expressed on a 14% grain moisture level, and grains smaller than 2.2 mm were sorted out and considered as fine grains.

For spring wheat, flag leaf measurements were taken on 23 July 2025. Six plants from each plot were randomly selected for flag leaf removal, and an additional six plants were chosen as control plants with intact flag leaves. At late wax maturity (BBCH 89), before mechanical harvesting, 12 marked plants (six with flag leaf removed and six control plants) were harvested manually from each plot for determination of plant yield components of the main shoot (grain number per spike and grain weight per spike). Spring wheat was harvested between 27 August and 8 September in 2025. Since spring wheat was a comparison with winter wheat in the flag leaf removal experiment, we did not measure final grain yield in spring wheat.

### 2.4. Stomatal anatomical analyses

For stomatal anatomical measurements, the central part of the flag leaf was used. Sampled leaf parts were cut in half widthwise (Fig. 2), and epidermal impressions of adaxial and abaxial surfaces were made using dental silicone (Speedex light body, Coltene/Whaledent AG for adaxial and oranwash L, Zhermack for abaxial). Then, nail varnish was applied to the sample impressions and transferred to microscope slides with transparent tape. An area of approximately 0.28 mm^2^ in the central part of the sample was captured using the Kern OBF 133 microscope and ODC 832 camera (Kern & Sohn GmbH) at 200x magnification. Stomatal density (SD, stomata mm^-2^), guard cell length (GCL, µm), guard cell width (GCW, µm) and pore length (PL, µm) were measured using ImageJ software (Schneider *et al*., 2012). For GCL, GCW and PL measurements, five stomata per sample were measured and average of these measurements was calculated to gain one representative value per plant. Total SD was calculated as the sum of adaxial and abaxial SDs. Stomatal density ratio (SR) was defined as the ratio of adaxial SD to abaxial SD (McAusland *et al*., 2021).

**Figure 2.**
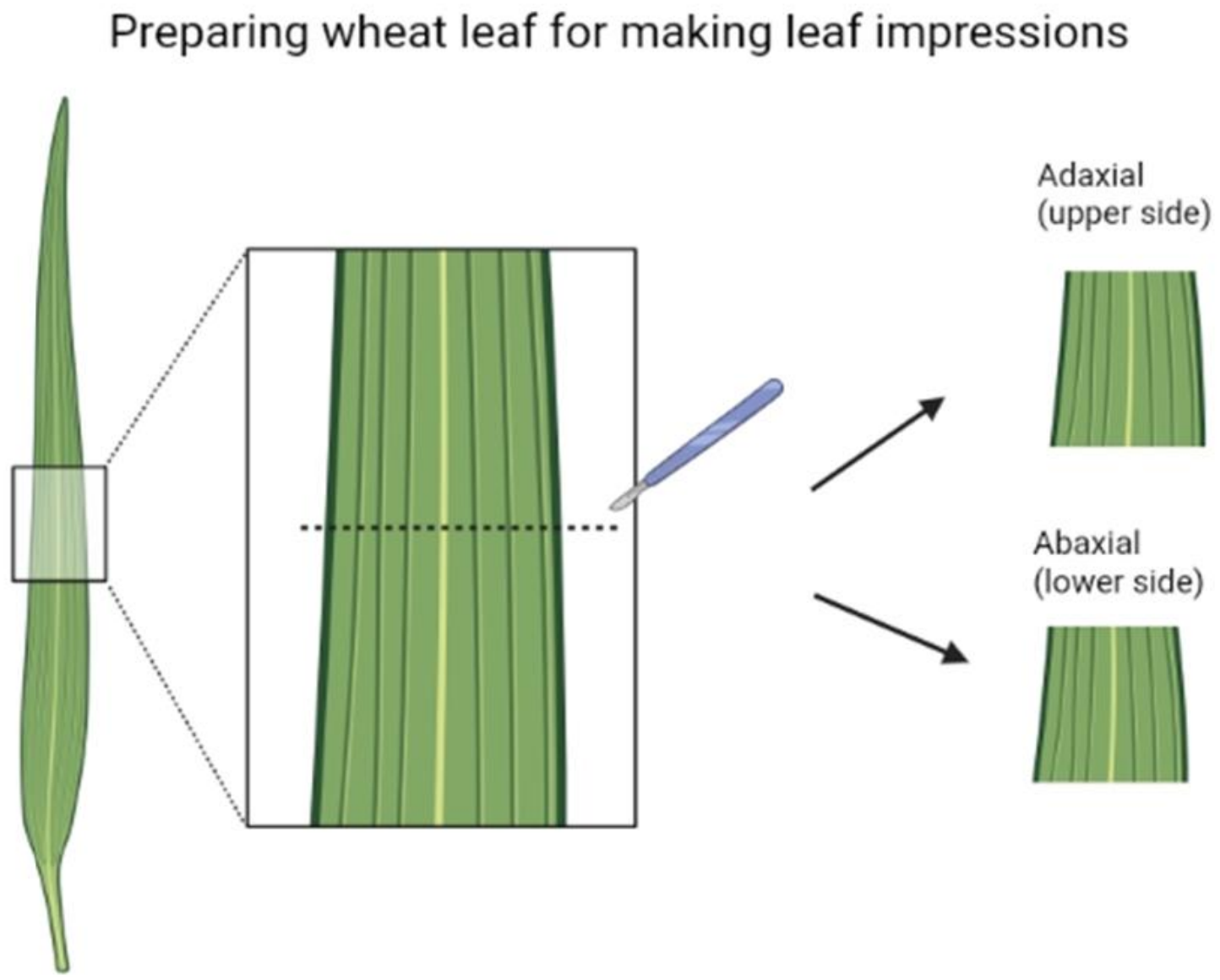
Figure illustrating the preparation of leaf impressions from a wheat flag leaf. First, we cut out the central part of the leaf. After that, we cut it in half crosswise from the centre and made epidermal impressions from the adaxial (upper) surface of one half and from the abaxial (lower) surface of the other half (figure created with BioRender.com) Alt text: Figure showing preparation of leaf impressions of wheat flag leaf.

Theoretical maximal anatomical stomatal conductance g_smax_ was calculated with the following equation (Caine *et al*., 2023):

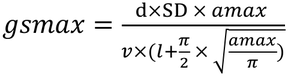

where d is diffusity of water in air at 25°C (0.0000249 m^-2^s^-1^) and *v* is molar volume of air (0.024464 m^3^ mol^-1^). SD is stomatal density, *amax* (µm^2^) is maximum pore area, calculated as abaxial or adaxial PL*PL/2, and pore depth *l* (µm) was considered to be equivalent to the width of guard cell at the centre of the stoma (Caine *et al*., 2023).

### 2.5. Agronomical traits and plant analyses

According to CPVO methodological guidelines (Community Plant Variety Office, 2019), plants were evaluated for glaucousness (GL) on a scale of 1–9 (1- absent or very weak, 9- very strong) about two weeks after heading. GL was evaluated on abaxial and adaxial flag leaf surfaces, and on spike.

Plant disease severity was assessed from the beginning of heading stage (BBCH 51) at approximately two-week intervals on three occasions during the growing season. Disease severity was rated on a scale of 1–9, where 1 indicated no visible signs of disease and 9 indicated that the entire leaf was infected. In 2024, conditions were favourable for the spread of plant diseases, and disease symptoms were visible even before heading. Therefore, assessments began earlier, at the end of stem elongation (BBCH 35), and were conducted on four occasions during the growing season. Plants were evaluated for the following diseases: yellow rust (*Puccinia striiformis*), tan spot (*Pyrenophora tritici-repentis*), powdery mildew (*Blumeria graminis*), and leaf blotch (*Zymoseptoria tritici*). The three-year cumulative disease index (CDI) was calculated as the summed incidence of all four diseases averaged over the three study years. The cumulative disease index for each year was calculated as the sum of disease incidences for all four diseases within one year. The disease index (DI) for each disease was calculated as the mean of its annual incidence values over the three years.

### 2.6. Statistical analyses

Analyses of variance (ANOVA) General Linear Models (GLM) procedure was used to evaluate differences in traits between genotypes and years. Homogeneity of variances was tested with the Levene’s test. Factorial analysis of variance (two-way ANOVA) was used to evaluate the main effects of genotype, year and their interactions for measured traits. Comparisons between individual means were performed using Tukey’s HSD post hoc test. Simple linear regression analyses were used to detect significant relationships between stomatal characteristics and grain yield. For nonparametric traits (disease index and glaucousness) the Kruskal-Wallis one-way ANOVA by ranks and multiple comparison of mean ranks were used. Spearman correlation was used for correlating nonparametric traits. For statistical analyses, measurements from six plants within each plot were averaged to obtain a single plot-level mean to be used in statistical analyses (n=3 plots per genotype within each year). For diseases, statistical analyses were performed using genotype-level means obtained by averaging values from 3 plots per genotype; for glaucousness, statistical analyses were performed using 3 plots per genotype within each year. For regression and correlation analyses, genotype-level means for each year were used. All effects were considered significant at p < 0.05. Statistical analyses were performed with Statistica (version 7.0, StatSoft Inc., Tulsa, OK, USA). All the graphs in the results section were made using GraphPad Prism, version 8.0 (GraphPad software, San Diego, California, USA, www.graphpad.com).

The manuscript was reviewed for language and clarity using Grammarly.

## 3. Results

### 3.1. Variation of agronomical traits of studied winter wheat varieties

#### 3.1.1. Grain yield

Grain yield was lowest in 2025 (average for early varieties 4140 kg ha^-1^ and late varieties 5097 kg ha^-1^), while in 2023 and 2024 it was higher, mean values for early varieties 7167 and 6149 kg ha^-1^ and for late varieties 8435 and 8878 kg ha^-1^, respectively (Fig. 3A,B). Grain yields of both early and late varieties were significantly affected by year, genotype, and their interaction (Table 1). The variability in grain yield was greater in 2024 than in 2023 and 2025 for early varieties, but more uniform in late varieties (Fig. 3A,B). In 2025, TGW was 1.5 times lower compared with 2023 in both early and late varieties (Fig. S1A,B). In 2024, Fredis and Ruske had the lowest TGW among early varieties (Fig. S1A). The proportion of fine grain in both early and late varieties was affected by genotype, year and their interaction (Table 1). In both early and late varieties, the proportion of fine grain was lowest in 2023 (mean 0.49 and 0.52%, respectively) and highest in 2025 (mean 16.3 and 13.7%, respectively) (Fig. S1C,D).

**Figure 3.**
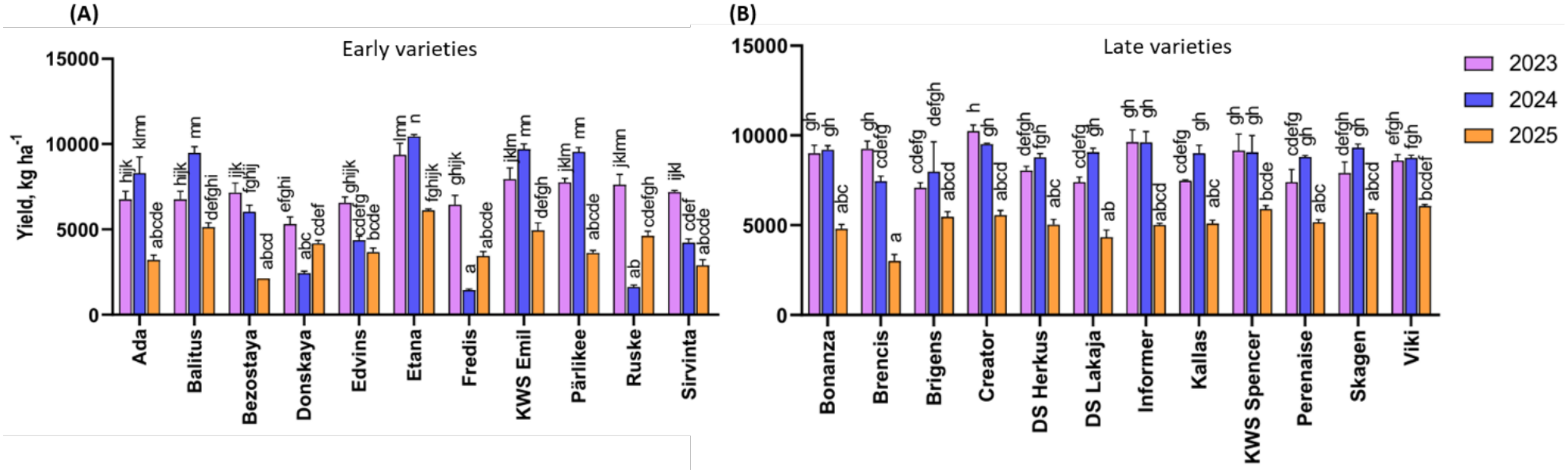
Grain yield (kg ha^-1^) of early and late varieties of winter wheat in 2023–2025; (A) Grain yield of early varieties in 2023–2025; (B) Grain yield of late varieties in 2023–2025. In 2025, two varieties (Dykanka and Sanzhara) were lodged in all replicates and, therefore, were not included in the harvest data. Error bars represent standard error of the mean and different letters indicate statistically significant differences between groups. Tukey’s post-hoc test was performed following two-way analysis of variance (ANOVA, p<0.05, n=3). Alt text: Graphs labelled A–B, showing winter wheat grain yield across three trial years in early and late varieties with statistical annotations.

**Table 1.**
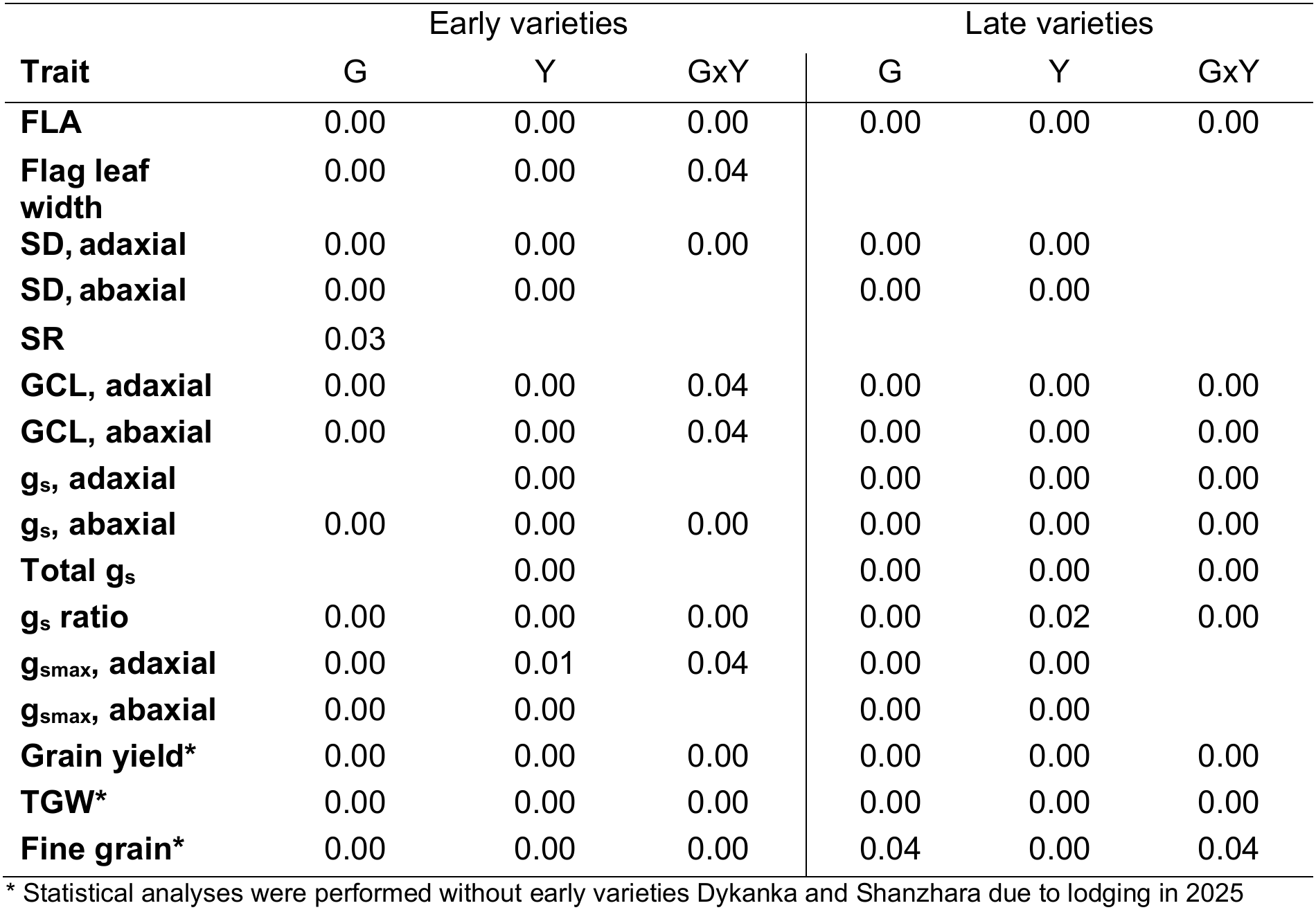
Results of factorial analyses of variance (Two-way ANOVA, p-values, n=3 plots per genotype per year) for genotype (G), year (Y), and GxY interaction in early and late varieties of winter wheat. Empty cells indicate no significant effects of studied factors (G, Y) or their interaction (GxY) on plant traits.

|  | Early varieties |  |  | Late varieties |  |  |
|---|---|---|---|---|---|---|
| **Trait** | G | Y | GxY | G | Y | GxY |
| **FLA** | 0.00 | 0.00 | 0.00 | 0.00 | 0.00 | 0.00 |
| **Flag leaf width** | 0.00 | 0.00 | 0.04 |  |  |  |
| **SD, adaxial** | 0.00 | 0.00 | 0.00 | 0.00 | 0.00 |  |
| **SD, abaxial** | 0.00 | 0.00 |  | 0.00 | 0.00 |  |
| **SR** | 0.03 |  |  |  |  |  |
| **GCL, adaxial** | 0.00 | 0.00 | 0.04 | 0.00 | 0.00 | 0.00 |
| **GCL, abaxial** | 0.00 | 0.00 | 0.04 | 0.00 | 0.00 | 0.00 |
| **g<sub>s</sub>, adaxial** |  | 0.00 |  | 0.00 | 0.00 | 0.00 |
| **g<sub>s</sub>, abaxial** | 0.00 | 0.00 | 0.00 | 0.00 | 0.00 | 0.00 |
| **Total g<sub>s</sub>** |  | 0.00 |  | 0.00 | 0.00 | 0.00 |
| **g<sub>s</sub> ratio** | 0.00 | 0.00 | 0.00 | 0.00 | 0.02 | 0.00 |
| **g<sub>smax</sub>, adaxial** | 0.00 | 0.01 | 0.04 | 0.00 | 0.00 |  |
| **g<sub>smax</sub>, abaxial** | 0.00 | 0.00 |  | 0.00 | 0.00 |  |
| **Grain yield\*** | 0.00 | 0.00 | 0.00 | 0.00 | 0.00 | 0.00 |
| **TGW\*** | 0.00 | 0.00 | 0.00 | 0.00 | 0.00 | 0.00 |
| **Fine grain\*** | 0.00 | 0.00 | 0.00 | 0.04 | 0.00 | 0.04 |

\* Statistical analyses were performed without early varieties Dykanka and Shanzhara due to lodging in 2025

In 2024, grain yield was remarkably low for early varieties Donskaya, Fredis, and Ruske (Fig. 3A), likely due to their susceptibility to yellow rust. We found no significant differences in cumulative disease index between years (Fig. S2A), but the variability in yellow rust infection scores in 2024 was high, likely due to the few susceptible early varieties (Fig. S2B). Cumulative disease index correlated negatively with grain yield in all individual years and in the pooled dataset of 2023–2025 (Fig. S3A,C−E). We also found a negative correlation between disease index of three diseases and grain yield (Fig. S3B).

#### 3.1.2. Glaucousness

Adaxial glaucousness (GL) was generally low, but slightly higher in 2024 and 2025 than in 2023 (Fig. S4). In warmer years (2023 and 2024), the abaxial GL was higher than in cooler year (2025) (Fig. S4). GL on the spike was highest in 2024 (Fig. S4). In the pooled dataset of 2023−2025, both abaxial and adaxial GL as well as GL on the spike were positively correlated with grain yield (Fig. S5A). Spike GL was significantly positively correlated with grain yield also in all individual years (Fig. S5A−D). Adaxial GL was positively correlated with grain yield in 2024 and 2025 (Fig. S5C,D), whereas abaxial GL was positively correlated with grain yield only in 2024 (Fig. S5C).

#### 3.1.3. Flag leaf removal

For both early and late varieties, genotype, year and their interaction had significant effects on flag leaf area (FLA), but only in early varieties was the flag leaf width affected by these factors (Table 1). An average FLA was nearly twice as large in 2025 as in 2023 (average for early varieties 33.95 and 16.82 cm^2^ and for late varieties 29.67 and 15.38 cm^2^, respectively) (Fig. S6A,B).

At the heading stage, we removed flag leaves from six randomly selected plants per plot to determine whether flag leaf removal affects grain yield components. Results showed a reduction in grain weight per spike of approximately 18% due to flag leaf removal in 2025 (Table 2), but there was no significant effect in 2023, 2024, nor in the pooled dataset (Table 2). We detected no effect of flag leaf removal on grain number per spike in any year in winter wheat.

**Table 2.**
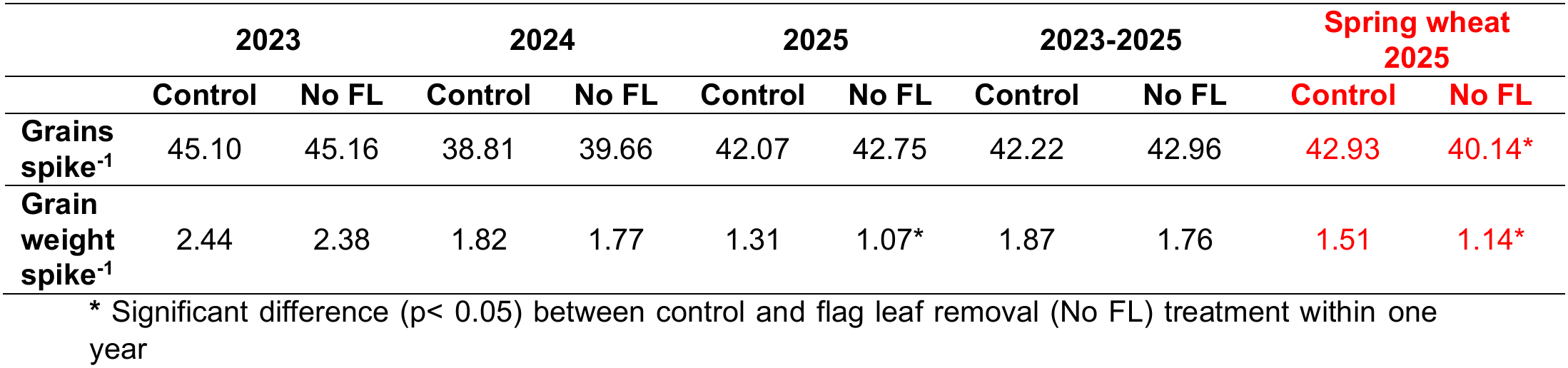
Factorial analysis of variance (Two-way ANOVA) of the effect of flag leaf (FL) removal on winter and spring wheat yield components (using genotype and treatment as factors, n=3 plots per genotype per year and n=23 varieties in the pooled dataset).

|  | **2023** |  | **2024** |  | **2025** |  | **2023-2025** |  | **Spring wheat 2025** |  |
|---|---|---|---|---|---|---|---|---|---|---|
|  | **Control** | **No FL** | **Control** | **No FL** | **Control** | **No FL** | **Control** | **No FL** | **Control** | **No FL** |
| **Grains spike<sup>-1</sup>** | 45.10 | 45.16 | 38.81 | 39.66 | 42.07 | 42.75 | 42.22 | 42.96 | 42.93 | 40.14* |
| **Grain weight spike<sup>-1</sup>** | 2.44 | 2.38 | 1.82 | 1.77 | 1.31 | 1.07* | 1.87 | 1.76 | 1.51 | 1.14* |

\* Significant difference (p< 0.05) between control and flag leaf removal (No FL) treatment within one year

In spring wheat in 2025, flag leaf removal had a stronger effect than in winter wheat: grain number per spike decreased by approximately 6.5%, and the grain weight per spike by 24% (Table 2).

### 3.2. Stomatal anatomical traits and their association with yield

Wheat considerably differs in stomatal density (SD) between leaf surfaces (p= 0.00, Table 3), with higher SD on the adaxial than on the abaxial surface (Fig. 4A-D). SD of both leaf surfaces was affected by genotype and year (Table 1); however, the GxY interaction was observed only for the adaxial surface of early varieties, where SDs were smaller in 2025 than in 2023 in most and smaller than 2024 in half of the studied varieties (Table 1, Fig. 4A-D). Despite large annual variation in SD, adaxial to abaxial stomatal density ratio (SR) was similar between years, and it was significantly affected by genotype only in early varieties (Table 1, Fig. 4E,F).

**Figure 4.**
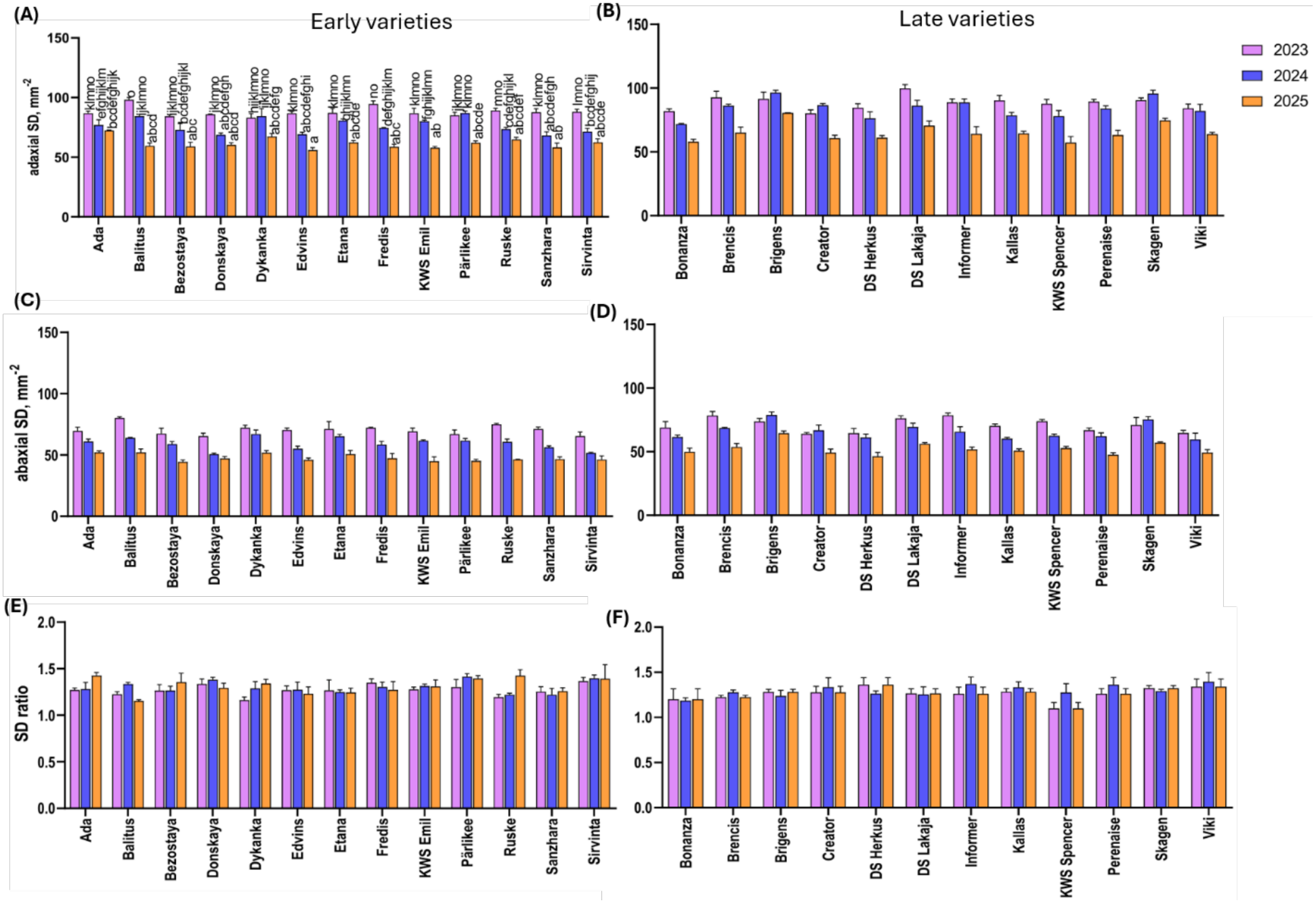
Stomatal densities (SD, stomata mm^-2^) and stomatal ratios of winter wheat flag leaves of early and late varieties in 2023−2025. (A) Adaxial SD of early varieties; (B) Adaxial SD of late varieties; (C) Abaxial SD of early varieties; (D) Abaxial SD of late varieties; (E) Stomatal ratio (SR, no unit) of early varieties; (F) Stomatal ratio of late varieties. Error bars represent standard errors of the means, and different letters indicate statistically significant differences between groups. Tukey’s post-hoc test was performed following two-way analysis of variance (ANOVA, p<0.05, n=3). Alt text: Graphs A−F, showing stomatal anatomical traits in 2023−2025. A is a bar chart of winter wheat adaxial stomatal density in early varieties. B is a bar chart of adaxial stomatal density in late varieties. C is a bar chart of abaxial stomatal density in early varieties. D is a bar chart of abaxial stomatal density in late varieties. E is a bar chart of adaxial to abaxial stomatal density ratio in early varieties. F is a bar chart of adaxial to abaxial stomatal density ratio in late varieties. All graphs are with statistical annotations.

**Table 3.** Results of factorial analyses of variance (Two-way ANOVA, n=25) of winter wheat flag leaf stomatal density (SD) for year, leaf surface, and year x leaf surface interaction on the abaxial and adaxial leaf surfaces of winter wheat.

|  | SS | Degr. of<br>Freedom | MS | F | p |
|---|---|---|---|---|---|
| Intercept | 710848.1 | 1 | 710848.1 | 19816.90 | 0.00 |
| year | 14424.2 | 2 | 7212.1 | 201.06 | 0.00 |
| leaf surface | 9568.6 | 1 | 9568.6 | 266.75 | 0.00 |
| Y x leaf surface | 158.1 | 2 | 79.1 | 2.20 | 0.11 |
| Error | 5165.4 | 144 | 35.9 |  |  |

Adaxial and abaxial GCLs were higher in 2025 compared with 2023 and 2024 for many early and few late varieties (Fig. S7A−D). We observed a positive relationship between adaxial SD and grain yield in 2024 and in the pooled dataset (Fig. 5A), and between abaxial SD and grain yield in 2024, 2025, and in the pooled dataset (Fig. 5B). We found a negative relationship between adaxial GCL and grain yield in two years out of three and in the pooled dataset (Fig. 5C), but abaxial GCL was negatively associated with grain yield only when the data from three years were pooled (Fig. 5D). SR was not significantly associated with grain yield in any individual years nor in the pooled data of 2023−2025 (Fig. 5E).

**Figure 5.**
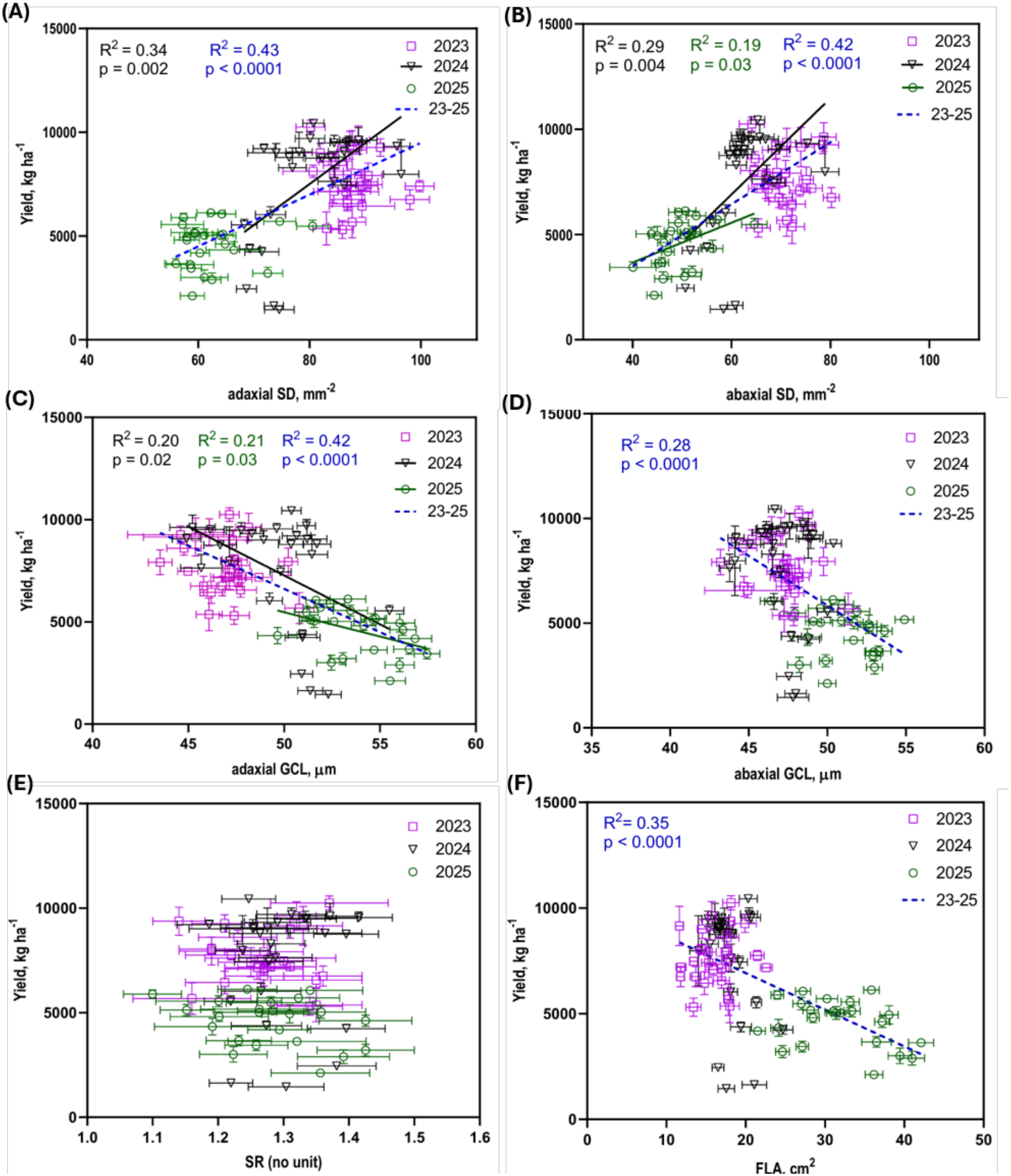
Linear regressions between grain yield and flag leaf stomatal traits of winter wheat. (A) Relationship between adaxial stomatal density (SD) and grain yield in 2023-2025; (B) Relationship between abaxial SD and grain yield in 2023−2025; (C) Relationship between adaxial guard cell length (GCL) and grain yield in 2023−2025; (D) Relationship between abaxial GCLand grain yield in 2023−2025; (E) Relationship between adaxial to abaxial stomatal density ratio (SR) and grain yield; (F) Relationship between flag leaf area (FLA) and grain yield. Graphs include linear regression lines for statistically significant relationships (p < 0.05, n= 23−25 genotypes within each year and 69−75 in the pooled dataset, error bars show standard errors of the means). Alt text: Graphs A−F, showing linear regressions between winter wheat grain yield and stomatal densities, stomatal size, stomatal ratio, and flag leaf area in individual years and in the pooled data. Stomatal traits and flag leaf area are on the x-axis and grain yield on the y-axis. All the graphs with significant relationships are with regression lines, statistical significances and p- values.

We found a negative relationship between FLA and grain yield in the pooled dataset of three years (Fig. 5F), and plants with larger flag leaves had lower SD on both leaf surfaces (Fig S8A,B). SD was negatively related with GCL in all individual years as well as in pooled data (Fig S8C,D). We also observed a negative correlation between both adaxial and abaxial SD and disease index for powdery mildew, and adaxial SD and disease index for yellow rust in the pooled dataset of 2023−2025 (Fig. S3B).

### 3.3. Anatomical maximal and operational stomatal conductance

To address maximum gas exchange potential of leaf surfaces, we calculated maximum anatomical stomatal conductance (g_smax_). Adaxial g_smax_ was affected by genotype, year and the GxY interaction in early varieties and by genotype and year in late varieties (Table 1), with lower values recorded in 2025 than in 2023 and 2024 in few early varieties (Fig. S9A-D). Abaxial g_smax_ was affected by genotype and year in both early and late varieties (Table 1). We observed a positive relationship between adaxial SD and adaxial g_smax_ in 2024 and 2025, and in the pooled dataset (Fig. S10A). Similar relationship for the abaxial surface was found in all individual years and in the pooled data of 2023−2025 (Fig. S10B). A negative relationship between g_smax_ and GCL for both leaf surfaces was evident only in the pooled data (Fig. S10C,D). We found positive relationships between adaxial and abaxial g_smax_ and grain yield in the pooled data, but not in individual years (Fig. S10E,F).

Operational adaxial g_s_ and total g_s_ of early varieties depended only on the year, but in other cases both the genotype, year and their interactions affected operational g_s_ and g_s_ ratio (Table 1). In 2025, adaxial g_s_ was higher than in 2023 in most and higher than in 2024 in some late varieties, whereas abaxial g_s_ was higher compared with 2023 in few varieties (Fig. S11A−D). The value of g_s_ ratio was over 2 (Fig. S11E,F), indicating that adaxial leaf surface is more important in gas exchange than abaxial surface. The g_s_ ratio of early varieties was more variable in 2025 compared with late varieties (Fig. S11E,F). Both early and late varieties exhibited the lowest abaxial and adaxial g_s_ /g_smax_ ratios in 2023 and the highest in 2025 (Fig. S12). We found a negative relationship between adaxial g_smax_ and operational g_s_ in the pooled dataset, but a positive relationship in 2024 (Fig. S13A), whereas abaxial g_smax_ was negatively related with operational g_s_ in the pooled dataset (Fig. S13B). We observed a positive relationship between adaxial g_s_ and grain yield only in 2023, whereas both adaxial and abaxial g_s_ and yield were negatively related in the pooled dataset (Fig. 6A,B). Similarly, the relationship between total g_s_ and grain yield was positive in 2023 and negative in the pooled dataset (Fig. 6C). Adaxial and abaxial g_s_ associated positively with each other in the pooled dataset but not in any individual years (Fig. 6D).

**Figure 6.**
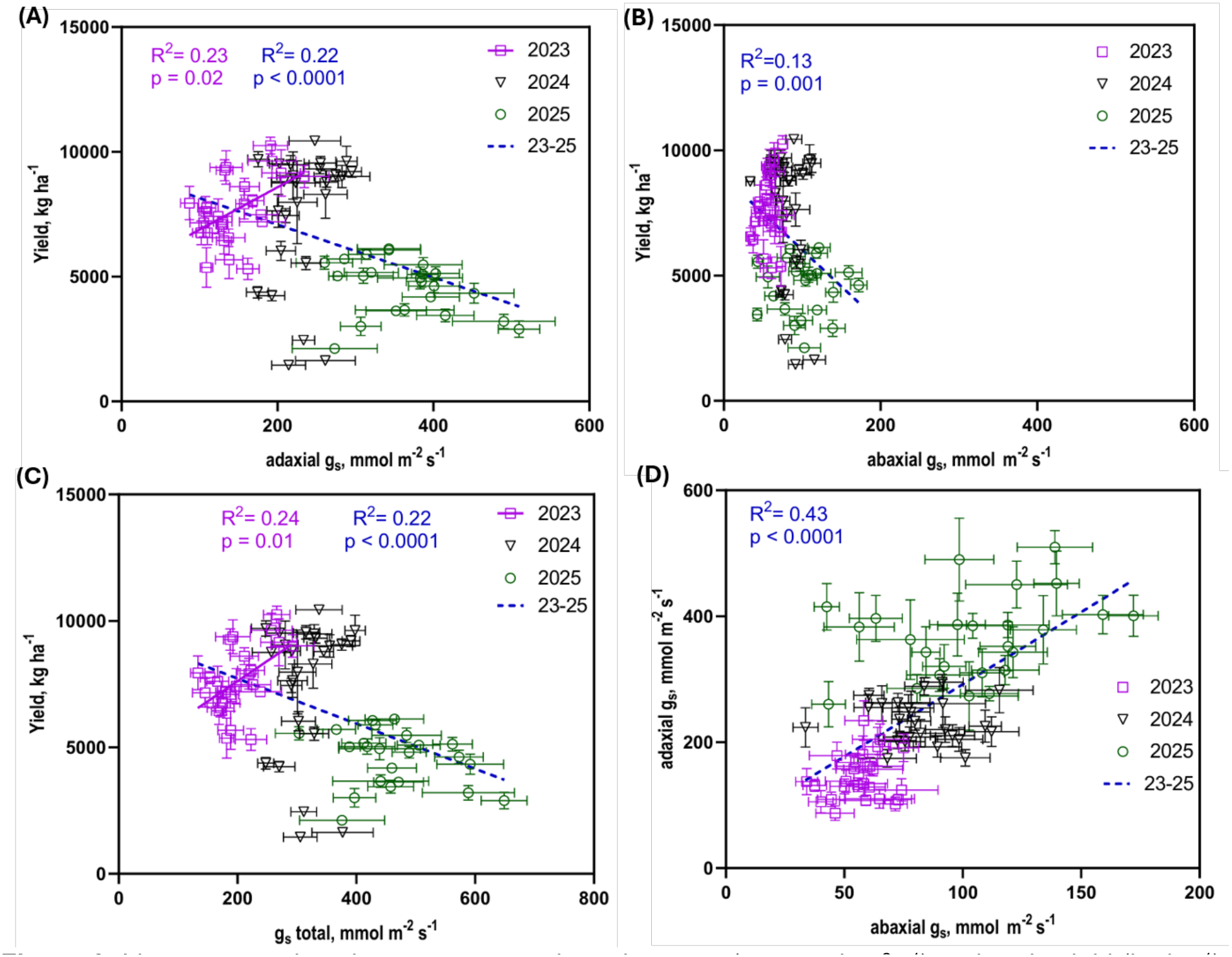
Linear regressions between stomatal conductance (g_s_, mmol m^-2^s^-1^) and grain yield (kg ha^-1^) on winter wheat. (A) Relationship between adaxial stomatal conductance and grain yield in 2023−2025; (B) Relationship between abaxial stomatal conductance and grain yield in 2023−2025; (C) Relationship between total stomatal conductance and grain yield; (D) Relationship between abaxial and adaxial stomatal conductance. Graphs include regression lines for statistically significant relationships (p < 0.05, n= 23−25 genotypes within each year and n=69−75 in the pooled dataset, error bars show standard error means). Alt text: Graphs labelled A−D. A is showing a linear relationship between winter wheat grain yield and adaxial stomatal conductance in individual years and in the pooled dataset of 2023−2025, B is showing a linear relationship between winter wheat grain yield and abaxial stomatal conductance in individual years and in the pooled dataset of 2023−2025, C is showing a linear relationship between winter wheat grain yield and total stomatal conductance in individual years and in the pooled dataset of 2023−2025, D is showing a linear relationship between abaxial and adaxial stomatal conductance in individual years and in the pooled dataset of 2023−2025. All the graphs with significant relationships are with regression lines, statistical significances and p-values.

## 4. Discussion

### 4.1. Winter wheat grain yield and effects of flag leaf removal on yield components

Precipitation and pathogens had major effects on yield in our study. Grain yield was lowest in 2025 due to heavy rains and lodging (Fig 1,3), in line with a previous report, where 10 mm increase in precipitation during the grain-filling period and close to maturity decreased winter wheat grain yield by 91−92 kg ha^-1^ (Peltonen-Sainio *et al*., 2011). Lodging occurred shortly after heading and affected several varieties, two of which could not be harvested. Such extensive lodging may have negatively affected grain filling by reducing leaf light exposure and canopy photosynthesis (Setter *et al*., 1997). Pathogens can account for 50−60% yield losses in susceptible varieties (Moldenhauer *et al*., 2006; Figueroa *et al*., 2018). Accordingly, disease severity was negatively correlated with yield in all years and in 2024, yellow rust was responsible for major yield losses in the few susceptible early varieties (Fig. 3, S2-3).

Flag leaf adaxial and abaxial and spike glaucousness (GL) were positively correlated with grain yield in the pooled dataset and often also in individual years (Fig. S5). Higher yields in glaucous plants irrespective of growth conditions have been found before (Johnson *et al*., 1983; Merah *et al*., 2000; Monneveux *et al*., 2004). Advantages of glaucousness may be associated with the ability of cuticular wax to lower leaf temperature via increased reflectance (Johnson *et al*., 1983), and reduce leaf water loss and increase water use efficiency (Richards *et al*., 1986). In case of spikes, high GL may lead to reduced ear wetting and less in-ear sprouting (King and von Wettstein-Knowles, 2000).

Flag leaf largely determines grain-filling rate, and may contribute up to 25% to final grain yield in spring wheat (Mahmood *et al*., 1991), so flag leaf removal at the end of the heading stage (BBCH 59) would be expected to lower grain yield components also in winter wheat. Contrary to our expectations, we found that removing the flag leaf significantly decreased grain weight per spike only in 2025 (Table 2) whereas no effect on grain number per spike was detected (Table 2). Other studies have reported a decrease in winter wheat grain weight per spike by ∼20-30%, in grain numbers per spike by 10−12% (Melahat Avci, 2005; Racz *et al*., 2022), and in grain yield by ∼12% (Patterson and Ohm, 1975). We also tested whether winter and spring wheat respond differently to flag leaf removal. Flag leaf removal may lower grain yield by up to 25% and TGW by up to 20%, whereas in some cultivars and studies it did not affect grain yield (Duwayri, 1984; Mahmood *et al*., 1991). Here, the flag leaf removal decreased spring wheat grain number per spike 6.5% and grain weight per spike 24%, which is more than in winter wheat (Table 2). These findings suggest that winter and spring wheat differ in their responses to flag leaf removal, highlighting differences in their physiology and in the contribution of the flag leaf to final grain yield. Winter wheat tends to have more shoots and greater leaf area (Watson *et al*., 1963; Lorenzo *et al*., 2015), and therefore can provide a larger residual photosynthetic source, which can help compensate for the loss of the flag leaf from the main shoot. Indeed, removing the flag leaf of the main shoot reduces the primary source of grain filling (Melahat Avci, 2005), but this deficiency may be compensated by the contributions of assimilates from other leaves (Song *et al*., 2025). Canopy shading may limit such compensation in cool and wet years that support the production of large foliage (Aggarwal *et al*., 1990), which might explain the reduction of grain yield components in winter wheat only in the cool and wet 2025 (Table 2). Spring wheat typically produces fewer tillers (Lorenzo *et al*., 2015), and therefore is more dependent on the main shoot flag leaf during grain filling. When the flag leaf in spring wheat is removed, the remaining tillers and leaves may be less able to compensate fully for the lost photosynthetic capacity, leading to a reduction in grain yield. Winter wheat may also accumulate more stem reserves during vegetative growth in spring-early summer compared with spring wheat, which has less time for foliage development. Stem reserves can then be used for grain growth and support higher individual grain weight and total yield (Dreccer *et al*., 2009). Considering the potential influence of weather conditions on the observed responses to flag leaf removal, further experiments across multiple environments are necessary to determine how weather conditions affect the impact of flag leaf removal on yield components and whether spring and winter wheat respond in a similar manner.

Unexpectedly, FLA was negatively associated with grain yield in the pooled dataset of three years, driven by data of 2025 (Fig. 5F). As fine grain proportion was particularly high in 2025 (Fig S1C,D), this relationship is likely explained by inability of plants to realise their yield potential due to cool and wet weather (Fig 1), as well as lodging, which likely hindered canopy photosynthesis. Although the flag leaf may contribute up to quarter of final grain yield, lodging- induced shading may have made actual canopy photosynthesis under field conditions, rather than the maximum photosynthetic capacity of individual leaves, more important for assimilate supply and grain filling as found in previous studies (Driever *et al*., 2014; Carmo-Silva *et al*., 2017).

### 4.2. Stomatal anatomical traits and their association with grain yield

We found higher adaxial than abaxial SD in winter wheat, resulting in SR values consistently above 1, in line with several previous works (Wang and Clarke, 1993; McAusland *et al*., 2021; Wall *et al*., 2022; Samantara *et al*., 2025). A recent study comparing wild (*T. dicoccoides*) and cultivated wheat (*T. durum*) (Cinti et al., 2026) found similarly higher adaxial than abaxial SD in cultivated but not in wild wheat, hinting at the possibility that higher adaxial SDs have been selected for in breeding. In eudicots, SR depends on growth environment, and increases under elevated vapour pressure deficit (VPD) in cotton (Devi and Reddy, 2018) and in *Arabidopsis thaliana* (Tulva *et al*., 2024). Although winter wheat SD was affected by both, genotype and year, SR was not affected by year (Table 1), indicating that SR was relatively stable across different environments. Similar stability of SR in different environments was found for rice (*O. sativa*) (Wang *et al*., 2023) and in CO_2_ elevation experiments for maize (*Zea mays*) (Driscoll *et al*., 2006). On the other hand, a significant GxY interaction effect on SR was found in spring wheat (Samantara *et al*., 2025), and few wheat varieties showed changes in SR when grown under elevated CO_2_ (Wall *et al*., 2023), suggesting environmental plasticity in SR in at least some wheat genotypes. Our findings suggest that in winter wheat, adjustments of stomatal density on the flag leaf surfaces in response to growth conditions are largely coordinated, leading to similar SRs even when SDs vary (Fig. 4A−E).

Having more stomata on the adaxial leaf surface may make the plant more susceptible to pathogens (Muir, 2015). All fungal pathogens evaluated in our study enter the plant through stomata (Schilder and Bergstrom, 1994; Moldenhauer *et al*., 2006; Solanki *et al*., 2019; Bernasconi *et al*., 2023; Rebaque *et al*., 2025), except *B. graminis,* which infects plant epidermal cells (Prats *et al*., 2007). Typically, higher SD is associated with increased susceptibility to pathogen invasion (Dutton *et al*., 2019; Tateda *et al*., 2019). We found no correlation between cumulative disease index and SD in the pooled dataset, but there was a negative correlation between adaxial SD and cumulative disease index in 2024 and abaxial SD and cumulative disease index in 2025 (Fig. S3D,E). We also observed no correlation between cumulative or individual disease indices and SR in any individual year nor in the pooled data (Fig. S3A-E), finding no evidence for the hypothesis that the cost of adaxial stomata is associated with susceptibility to pathogens (Muir, 2015, 2020). As variation of SR in our study was minor (Fig. 4), the effects of stomatal traits may have been outweighted by other mechanisms influencing disease susceptibility.

Higher SD is often associated with smaller stomata (Hetherington and Woodward, 2003; Drake *et al*., 2013) and in our dataset, SD was also negatively associated with GCL (Fig. S8C,D). Both small stomatal size and high SR are associated with faster stomatal responses (Hetherington and Woodward, 2003; Franks and Farquhar, 2007; Drake *et al*., 2013; Woning *et al*., 2026; Yang *et al*., 2026), potentially allowing for increased sensitivity of gas-exchange to abiotic environment. For example, rice plants with smaller stomata and higher SD closed their stomata faster in response to rising temperature and VPD, thereby improving water use efficiency (WUE) (Caine *et al*., 2023). SD together with GCL and PL determine maximal potential for gas exchange (g_smax_). Our study showed that although g_smax_ in both early and late varieties was higher in warm 2023 compared with cooler 2025 (Fig. S9A-D), driven by higher SDs in 2023 (Fig. 4, S10), operational g_s_ values were lowest in 2023 and highest in 2025 (Fig. S11A-D), leading to a negative relationship between g_smax_ and g_s_ in the pooled dataset (Fig. S13. Similar decoupling between anatomical capacity for gas-exchange and realised conductance was also found in spring wheat, indicating strong plasticity of operational g_s_ (Chaplin *et al*., 2026b). We also found that abaxial g_smax_ was positively related with SD in all years and adaxial g_smax_ in 2024 and 2025 (Fig. S10A,B), whereas negative correlation between g_smax_ and GCL was found only in the pooled data (Fig. S10C,D). These results are in line with previous findings suggesting that SD and not stomatal size is the primary factor affecting g_smax_ (Ochoa et al., 2024; Wall et al., 2022).

Stomatal development in grasses occurs in distinct rows along veins, with the earliest stages occurring basally, and during the development, cells move upwards as the leaf expands longitudinally (Hepworth *et al*., 2018; Nunes *et al*., 2020), leading to larger but less dense stomata during flag leaf expansion (Zanella *et al*., 2023). In line with this, we observed a negative correlation between SD and FLA on both leaf surfaces (Fig. S8A,B). A similar relationship has been noted before in winter wheat (Morgan and LeCain, 1991), and also in e.g. eucalyptus (Franks *et al*., 2009), mangroves (Peel *et al*., 2017), barley (Samantara *et al*., 2026), and spring wheat (Samantara et al., 2025), and is line with adaptation of cell expansion growth to environmental conditions, leading to altered stomatal spacing (Granier and Tardieu, 2009; Tulva *et al*., 2024).

We found a positive association between both adaxial and abaxial SD and grain yield in the pooled data and in some individual years (Fig. 5A,B). Similarly, other studies with winter wheat have found a positive relationship between abaxial SD and grain yield (Yang *et al*., 2022; Du *et al*., 2026) and between adaxial SD and grain yield (Baruah *et al*., 2017). However, in spring wheat, a negative relationship between adaxial but not abaxial SD and grain yield was found in warm years 2022 and 2023 and was interpreted as costly water loss via adaxial stomata (Samantara *et al*., 2025). Our study included two warm years 2023 and 2024, and relatively cool, wet 2025. Even in 2023, we found no negative association between adaxial SD and grain yield, perhaps due to the different timing of development in spring and winter wheat that may have made winter wheat less susceptible to the dry spring and early summer of 2023. Spring and winter wheat may respond differently to contrasting climate conditions, as demonstrated also by the effects of flag leaf removal on yield components in winter and spring wheat in our experiment (Table 2). We found no significant relationship between SD ratio and grain yield in any individual year or in the pooled data, perhaps due to the minor variation in SR between years (Fig. 4, 5E). Stomatal size on both leaf surfaces was negatively associated with grain yield in the pooled data (Fig 5C,D), in line with another recent wheat field study (Chaplin *et al*., 2026b). In two out of three individual years, adaxial stomatal size also negatively associated with grain yield (Fig. 5C,D), perhaps suggesting that smaller, potentially faster adaxial stomata are particularly important for plasticity of flag leaf gas-exchange. Indeed, plasticity of adaxial stomatal traits was suggested to be particularly important for wheat adaptation to growth environment in the field (Chaplin *et al*., 2026b). While some studies suggest that smaller stomata are not necessarily faster in wheat (Woning *et al*., 2026) or across species (Elliott-Kingston *et al*., 2016), together, our findings support the hypothesis that smaller, denser stomata that allow for faster responses and higher WUE (Hetherington and Woodward, 2003; Franks and Farquhar, 2007; Drake *et al*., 2013; Caine *et al*., 2023; Yoshiyama *et al*., 2024), are favourable under variable field conditions.

### 4.3. Adaxial and abaxial stomatal conductance

Stomatal conductance is dependent on stomatal density, size and aperture, and is an important component of leaf CO_2_ conductance, whereas surface-specific differences in conductance may lead to different contribution of leaf surfaces to photosyhthesis (Muir, 2015; Ouyang *et al*., 2017; Wall *et al*., 2022; Ochoa *et al*., 2024). We observed a positive linear relationship between adaxial and abaxial g_s_ in the pooled data but not in any individual years (Fig. 6D). The absence of significant relationship within individual years suggests that stomatal responses to environmental factors are independent in the two leaf surfaces in short timescales, but coordination of g_s_ between surfaces is evident in the broad range of g_s_ values measured across the years. Independent stomatal responses between leaf surfaces were found before in wheat, whereas adaxial stomata responded faster to changes in light intensity than abaxial stomata (Wall et al., 2022). Alternatively, our results may be explained by differences in abiotic conditions between leaf surfaces at measurement time.

Stomatal conductance often positively correlates with grain yield (Fischer *et al*., 1998; Aminian *et al*., 2011; Roche, 2015). In our study, adaxial and total g_s_ were positively associated with grain yield only in 2023 (Fig. 6A,C), confirming that in warm and dry years, maintaining higher gas-exchange may be beneficial for yield in Nordic conditions (Jalakas *et al*., 2018). In the pooled dataset, the relationships between g_s_ and grain yield were negative (Fig. 6A-C). These results were driven by the data of the cool, wet 2025, where grain yield was low due to lodging (Fig. 1,3). Single timepoint measurements of g_s_ in June were likely not sufficient to predict the contribution of stomata to grain yield throughout the growing season. Stomatal anatomical traits may better explain variation in grain yield, as they capture the environmental variation throughout plant growth and development in the field.

## 5. Conclusions

Contrary to expectations, flag leaf removal in winter wheat had little effect on grain yield components, which is different from spring wheat and winter wheat grown in warmer climates. Thus, in winter wheat, the function of the flag leaf may be compensated by other leaves, as winter wheat usually has more tillers than spring wheat, and by longer time to accumulate stem reserves for further translocation to grains.

The major aim of this study was to examine winter wheat stomatal traits and their relationships with grain yield under field conditions across three years. We found that in warm, dry years, plants produce smaller leaves with higher SD. Despite significant variation in growth environment and SD, SR remained stable, indicating that wheat plants increase or decrease the number of stomata proportionally on leaf surfaces. At the same time, stomata on both leaf surfaces may respond differently to environmental factors, as we observed no correlation between adaxial and abaxial g_s_ values in individual years.

Higher stomatal densities and smaller stomatal sizes are associated with higher grain yield, potentially reflecting the benefit of faster stomatal responsiveness achieved by smaller stomata. Correlations between g_s_ and grain yield were inconsistent, highlighting the variability and plasticity of g_s_. Varieties with higher cumulative disease scores had lower grain yield as expected, but higher SD was not accompanied by stronger infection by stomata-entering pathogens, suggesting that traits other than SD largely determined disease susceptibility in the field.

Our results provide insight into variation of leaf-surface specific stomatal traits under field conditions in winter wheat, and on their associations with grain yield. Our findings suggest that breeding towards varieties with smaller, faster stomata at high densities may be beneficial for wheat yield under field conditions.

## Abbreviations

FL: flag leaf
FLA: flag leaf area
GCL: guard cell length
GL: glaucousness
g_s_: stomatal conductance
g_smax_: anatomical maximum stomatal conductance
PL: pore length
SD: stomatal density
SR: adaxial to abaxial stomatal density ratio
TGW: thousand grain weight

## Acknowledgements

We thank Mikk Välbe for helping us with sample collection and support in laboratory. We are also grateful to Made Ratassepp, Külli Jõessar, Jane Suvvi, Veste Vapper, and Merilin Saar for the help with field works.

## Authors contribution

H.H., E.I., E.M. and R.K. conceived and designed the study; E.I., P.J., I.T., M.K., K.S., B.K.D., A.I., M.H., R.K., E.M., and H.H. collected and analysed data; E.I and H.H wrote the manuscript with input from all authors.

## Data availability

All primary data to support the findings of this study are available in Supplementary Table S2 and will be deposited in a data repository upon acceptance of the manuscript.

