## Supplementary material for "Higher stomatal density and smaller stomatal size associate with higher grain yield in winter wheat"

**Table S1.** Winter wheat varieties used in field trials in 2023–2025.

| Variety | Type | Early/ Late variety | Origin country | Registration in origin country[1] | Registration in Estonia[2] |
|---|---|---|---|---|---|
| Ada | Winter | Early | Lithuania | 2004 | 2005 |
| Balitus | Winter | Early | Germany | 2015 | 2019 |
| Donskaya-Polukarlikovaya | Winter | Early | Russia, Rostov Region | *No data* | |
| Edvins | Winter | Early | Latvia | 2012 | 2013 |
| Fredis | Winter | Early | Latvia | 2008 | 2011 |
| Ruske | Winter | Early | Estonia | 2017 | 2017 |
| Etana | Winter | Early | Germany | 2013 | 2015 |
| KWS Emil | Winter | Early | Germany | 2016 | 2017 |
| Dykanka | Winter | Early | Ukraine | 2007[3] | |
| Sanzhara | Winter | Early | Ukraine | 2017[4] | |
| Pärlikee | Winter | Early | Estonia | 2025 | 2025 |
| Bezostaya | Winter | Early | Russia | 1959[5] | |
| Širvinta 1 | Winter | Early | Lithuania | 1989[6] | 1991[7] |
| Bonanza | Winter | Late | Germany | 2015 | 2017 |
| Creator | Winter | Late | Denmark | 2014 | 2016 |
| Kallas | Winter | Late | Estonia/ Lithuania | 2013 | 2013 |
| KWS Spencer | Winter | Late | Germany | 2017 | 2019 |
| Perenaise | Winter | Late | Estonia | 2019 | 2019 |
| Skagen | Winter | Late | Sweden | 2006 | 2011 |
| DS Herkus | Winter | Late | Lithuania | 2016 | 2016 |
| DS Lakaja | Winter | Late | Lithuania | 2019 | 2020 |
| Brigens | Winter | Late | Latvia | 2022 | |
| Brencis | Winter | Late | Latvia | 2017 | 2019 |
| Informer | Winter | Late | Germany | 2018 | |
| Viki | Winter | Late | Czech | 2018 | |

[1] Source: Community Plant Verify Office (CPVO) variety finder (https://cpvo.europa.eu/en/applications-and-examinations/cpvo-variety-finder )

[2] Source: Estonian plant variety register (https://portaal.agri.ee/avalik/#/sort)

[3] Source: https://sort.sops.gov.ua/cultivar/view/11252

[4] Source: https://agronom.info/en/p/wheat-sanzhara-37169.html

[5] Source: doi:10.1017/S0021859610000146

[6] Source: https://www.vle.lt/straipsnis/sirvinta-1/

[7] Source: https://www.riigiteataja.ee/et/akt/81819?leiaKehtiv

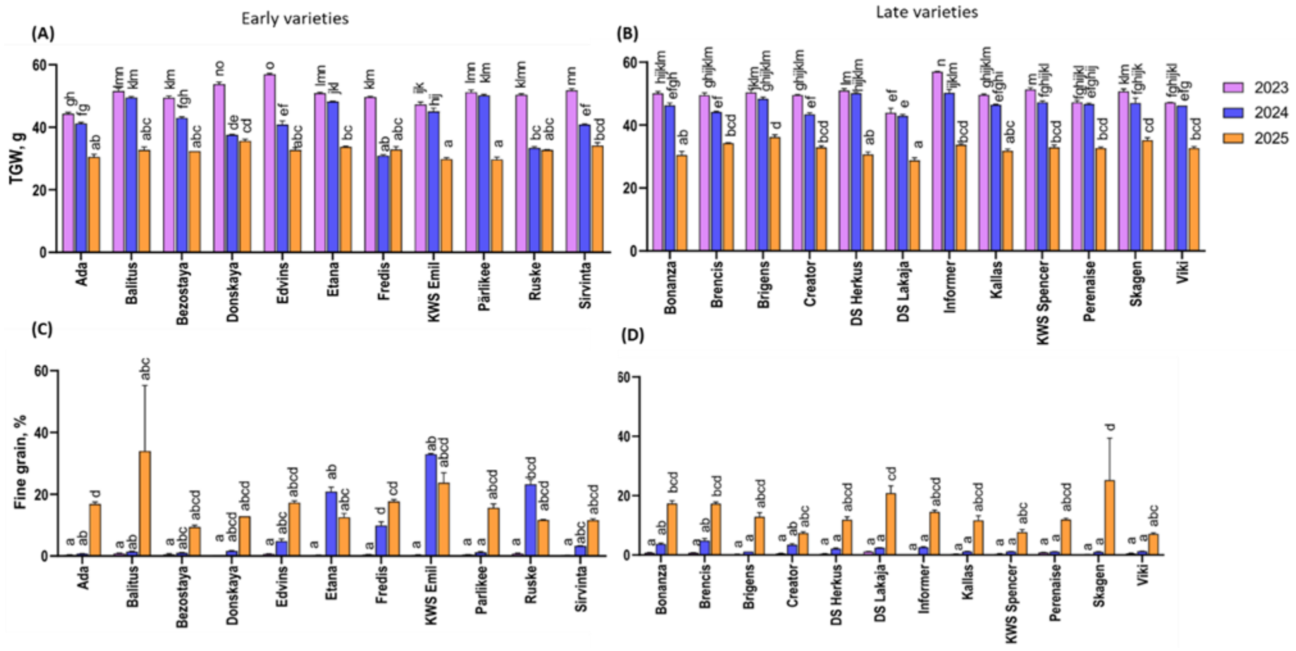


**Figure S1.** Thousand grain weight (TGW, g) and the proportion of fine grain (%) during three trial years in early and late varieties of winter wheat. (A) TGW of early varieties in 2023–2025; (B) TGW of late varieties in 2023–2025; (C) Proportion of fine grain of early varieties in 2023–2025; (D) Proportion of fine grain in 2023–2025. In 2025, two early varieties (Dykanka and Sanzhara) were lodged in all replicates and were therefore not included in the harvest data. Error bars represent standard errors of the means, and different letters indicate statistically significant differences between groups. Tukey's post-hoc test following factorial analysis of variance (two-way ANOVA, $p < 0.05$, $n=3$) was used.

Alt text: Graphs A–B, showing winter wheat thousand grain weight and the proportion of fine grain in early and late varieties in 2023–2025 with statistical annotations.

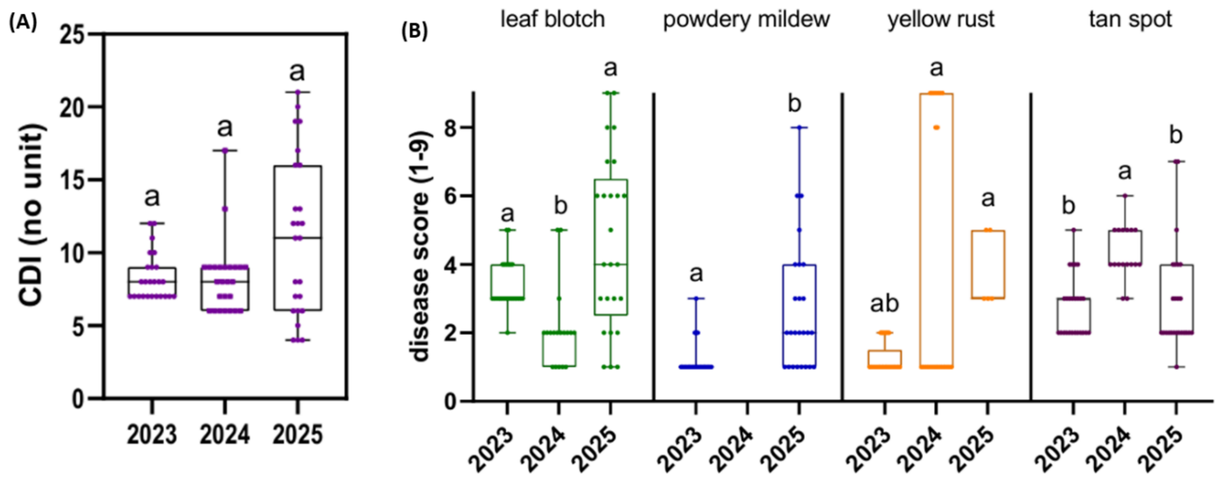


**Figure S2**. Fungal diseases in 2023–2025. (A) Cumulative disease index (CDI, no unit) in winter wheat in 2023–2025; Box-plot whiskers indicate minimum and maximum values and horizontal line in the box represents the median; (B) Winter wheat plant diseases scores on a scale 1–9 (1 indicates no visible signs of disease and 9 indicates that the entire leaf is infected) in 2023–2025. Letters indicate statistically significant differences between years within one disease (Kruskal-Wallis test, $p < 0.05$, n= 25).

Alt text: Graphs A–B. A is showing winter wheat cumulative disease index in three trial years. On the x-axis are study years and on the y-axis is cumulative disease index (CDI). B is showing winter wheat plant diseases scores in 2023–2025. On the x-axis are study years and on the y-axis are disease scores 1–9. Graphs are with statistical annotations.

(A) 2023-2025

| | CDI | yield | SD_ab | SD_ad | SD ratio |
|---|---|---|---|---|---|
| CDI | 1 | | | | |
| yield | -0.84 | 1 | | | |
| SD_ab | | | 1 | | |
| SD_ad | | | 0.79 | 1 | |
| SD ratio | | | -0.58 | | 1 |

(B) 2023-2025

| | DI_LB | DI_PM | DI_YR | DI_TS | yield | SD_ab | SD_ad | SD ratio |
|---|---|---|---|---|---|---|---|---|
| DI_LB | 1 | | | | | | | |
| DI_PM | 0.61 | 1 | | | | | | |
| DI_YR | 0.52 | 0.68 | 1 | | | | | |
| DI_TS | 0.40 | | | 1 | | | | |
| yield | -0.70 | -0.67 | -0.84 | | 1 | | | |
| SD_ab | | -0.59 | | | | 1 | | |
| SD_ad | | -0.48 | -0.42 | | | 0.79 | 1 | |
| SD ratio | | | | | | -0.58 | | 1 |

(C) 2023

| | CDI | yield | SD_ab | SD_ad | SD ratio |
|---|---|---|---|---|---|
| CDI | 1 | | | | |
| yield | -0.60 | 1 | | | |
| SD_ab | | | 1 | | |
| SD_ad | | | 0.69 | 1 | |
| SD ratio | | | -0.68 | | 1 |

(D) 2024

| | CDI | yield | SD_ab | SD_ad | SD ratio |
|---|---|---|---|---|---|
| CDI | 1 | | | | |
| yield | -0.52 | 1 | | | |
| SD_ab | | 0.58 | 1 | | |
| SD_ad | -0.42 | 0.58 | 0.88 | 1 | |
| SD ratio | | | | | 1 |

(E) 2025

| | CDI | yield | SD_ab | SD_ad | SD ratio |
|---|---|---|---|---|---|
| CDI | 1 | | | | |
| yield | -0.57 | 1 | | | |
| SD_ab | -0.42 | 0.47 | 1 | | |
| SD_ad | | | 0.6 | 1 | |
| SD ratio | | | | 0.47 | 1 |

**Figure S3.** Spearman correlation matrix of cumulative disease index (CDI) and single disease indices (DI) with grain yield and stomatal traits. (A) Spearman correlation matrix between cumulative disease index (CDI) and grain yield and stomatal densities (SD); (B) Spearman correlation matrix between disease index (DI) in 2023-2025 (LB- leaf blotch, PM- powdery mildew, YR- yellow rust, TS- tan spot); (C–E) Spearman correlation matrix between cumulative disease index (CDI) and stomatal traits and grain yield in 2023, 2024, and 2025, respectively; ($R_{Spearman}$, $p < 0.05$, N=25). Positive correlations are represented in red and negative in blue, empty cells indicate non-significant correlations.

Alt text: Graphs A–E, showing Spearman correlations between winter wheat cumulative disease index (CDI) and grain yield, and between disease indices (DI) and grain yield in three trial years. Statistically significant correlations are marked in bold.

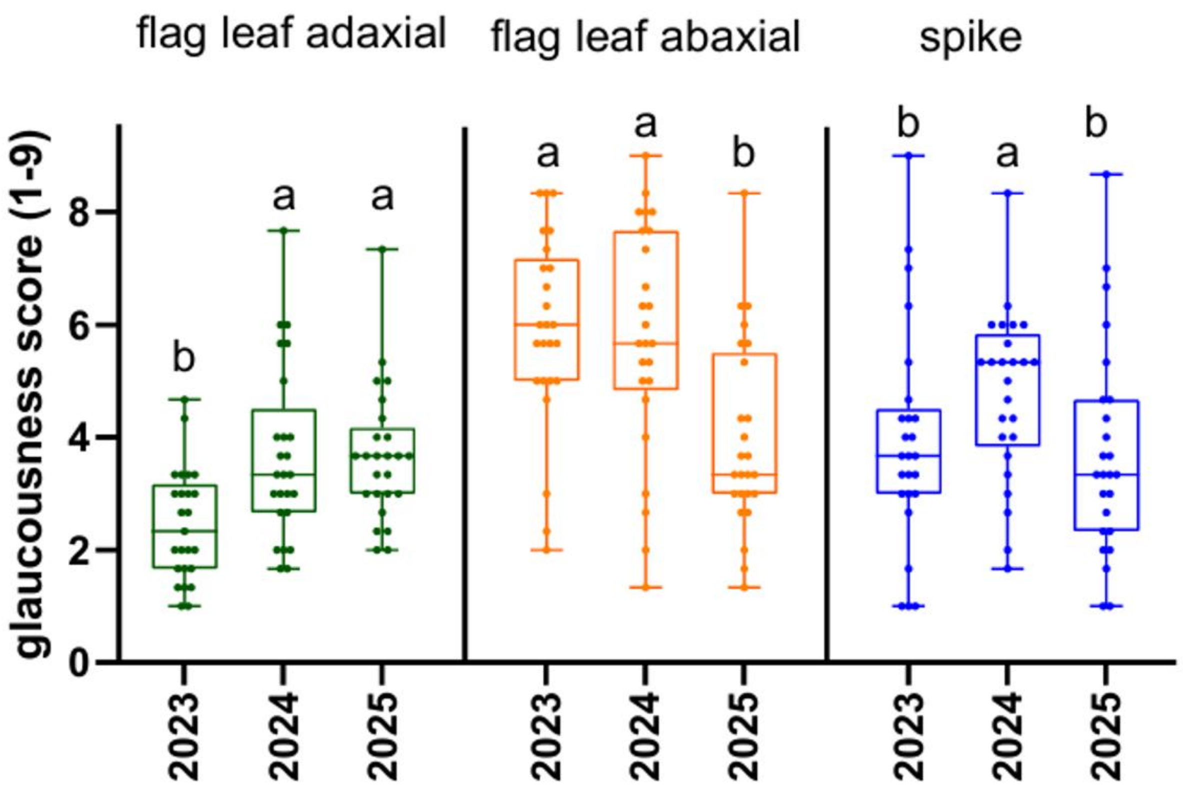


**Figure S4.** Glaucousness (GL) scores (1–9, where 1 is weak or absent and 9 is very strong) across the trial years of 2023-2025. Whiskers represent minimum and maximum values and horizontal line in the box represents the median. Letters indicate statistically significant differences between years within one plant organ (Kruskal-Wallis test, $p < 0.05$, n= 25).

Alt text: Graph showing winter wheat glaucousness scores (1–9, where 1 is weak or absent and 9 is very strong) across trial years of 2023–2025 with statistical annotations.

(A) 2023-2025

| | GL_ab | GL_ad | GL_spike | yield | SD_ab | SD_ad | $g_s$_ab | $g_s$_ad | $g_{smax}$_ab | $g_{smax}$_ad |
|---|---|---|---|---|---|---|---|---|---|---|
| GL_ab | 1 | | | | | | | | | |
| GL_ad | 0.88 | 1 | | | | | | | | |
| GL_spike | 0.74 | 0.8 | 1 | | | | | | | |
| yield | 0.66 | 0.64 | 0.79 | 1 | | | | | | |
| SD_ab | | | | | 1 | | | | | |
| SD_ad | | | | | 0.81 | 1 | | | | |
| $g_s$_ab | 0.47 | 0.45 | | | | | 1 | | | |
| $g_s$_ad | | | | | | | | 1 | | |
| $g_{smax}$_ab | | | | | 0.55 | 0.5 | | | 1 | |
| $g_{smax}$_ad | | | | | 0.58 | 0.54 | | | 0.87 | 1 |

(B) 2023

| | GL_ab | GL_ad | GL_spike | yield | SD_ab | SD_ad | $g_s$_ab | $g_s$_ad | $g_{smax}$_ab | $g_{smax}$_ad |
|---|---|---|---|---|---|---|---|---|---|---|
| GL_ab | 1 | | | | | | | | | |
| GL_ad | 0.73 | 1 | | | | | | | | |
| GL_spike | 0.51 | 0.49 | 1 | | | | | | | |
| yield | | | 0.74 | 1 | | | | | | |
| SD_ab | | | | | 1 | | | | | |
| SD_ad | | | | | 0.69 | 1 | | | | |
| $g_s$_ab | 0.49 | | | | | | 1 | | | |
| $g_s$_ad | | | | 0.43 | | | | 1 | | |
| $g_{smax}$_ab | | | | | 0.56 | 0.4 | 0.44 | | 1 | |
| $g_{smax}$_ad | | | | | | 0.41 | | -0.4 | 0.7 | 1 |

(C) 2024

| | GL_ab | GL_ad | GL_spike | yield | SD_ab | SD_ad | $g_s$_ab | $g_s$_ad | $g_{smax}$_ab | $g_{smax}$_ad |
|---|---|---|---|---|---|---|---|---|---|---|
| GL_ab | 1 | | | | | | | | | |
| GL_ad | 0.88 | 1 | | | | | | | | |
| GL_spike | 0.76 | 0.85 | 1 | | | | | | | |
| yield | 0.79 | 0.64 | 0.58 | 1 | | | | | | |
| SD_ab | 0.47 | | | 0.58 | 1 | | | | | |
| SD_ad | 0.47 | | | 0.58 | 0.88 | 1 | | | | |
| $g_s$_ab | | | | | | | 1 | | | |
| $g_s$_ad | | | | | | | | 1 | | |
| $g_{smax}$_ab | | | | | 0.58 | 0.48 | | | 1 | |
| $g_{smax}$_ad | | | | | 0.51 | 0.55 | | | 0.76 | 1 |

(D) 2025

| | GL_ab | GL_ad | GL_spike | yield | SD_ab | SD_ad | $g_s$_ab | $g_s$_ad | $g_{smax}$_ab | $g_{smax}$_ad |
|---|---|---|---|---|---|---|---|---|---|---|
| GL_ab | 1 | | | | | | | | | |
| GL_ad | 0.81 | 1 | | | | | | | | |
| GL_spike | 0.72 | 0.79 | 1 | | | | | | | |
| yield | | 0.44 | 0.45 | 1 | | | | | | |
| SD_ab | | | | 0.47 | 1 | | | | | |
| SD_ad | | | | | 0.6 | 1 | | | | |
| $g_s$_ab | 0.51 | | | | | | 1 | | | |
| $g_s$_ad | | | -0.58 | | | | | 1 | | |
| $g_{smax}$_ab | | | | | 0.55 | | | | 1 | |
| $g_{smax}$_ad | | -0.5 | | | | 0.54 | | | | 1 |

**Figure S5.** Spearman correlations between glaucousness (GL) of flag leaf surfaces and spike, grain yield, and stomatal traits in winter wheat. (A) Correlations between GL, grain yield and stomatal traits as average of 2023−2025; (B) Correlations between GL, grain yield and stomatal traits in 2023; (C) Correlations between GL, grain yield and stomatal traits in 2024; (D) Correlations between GL, grain yield and stomatal traits in 2025. ($R_{Spearman}$, $p < 0.05$, N=25). Positive correlations are represented in red and negative in blue, empty cells indicate non-significant correlations.

Alt text: Graphs A−D, showing Spearman correlations between winter wheat glaucousness of flag leaf surfaces and spike, grain yield, and stomatal traits in 2023–2025. Statistically significant correlations are marked in bold.

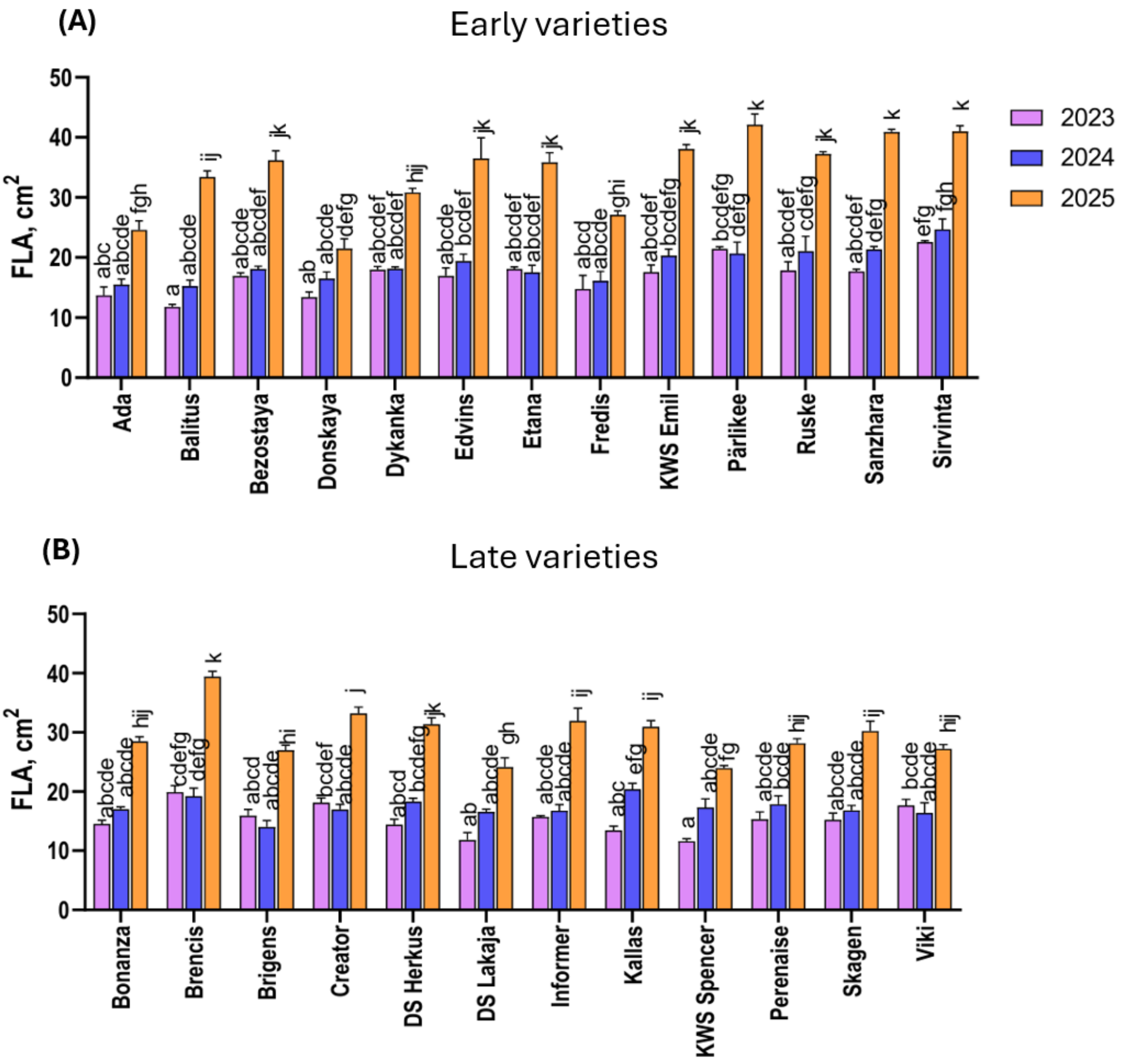


**Figure S6.** Flag leaf area (FLA, $cm^2$) of early and late winter wheat varieties in 2023–2025. (A) FLA of early varieties in 2023–2025; (B) FLA of late varieties in 2023–2025. Error bars represent standard error of the mean, and different letters indicate statistically significant differences between groups. Tukey's post-hoc test following factorial analysis of variance (two-way ANOVA, $p < 0.05$, $n=3$) was used.

Alt text: Graphs A and B, showing winter wheat flag leaf area (FLA, $cm^2$) of early and late varieties in 2023–2025 with statistical annotations.

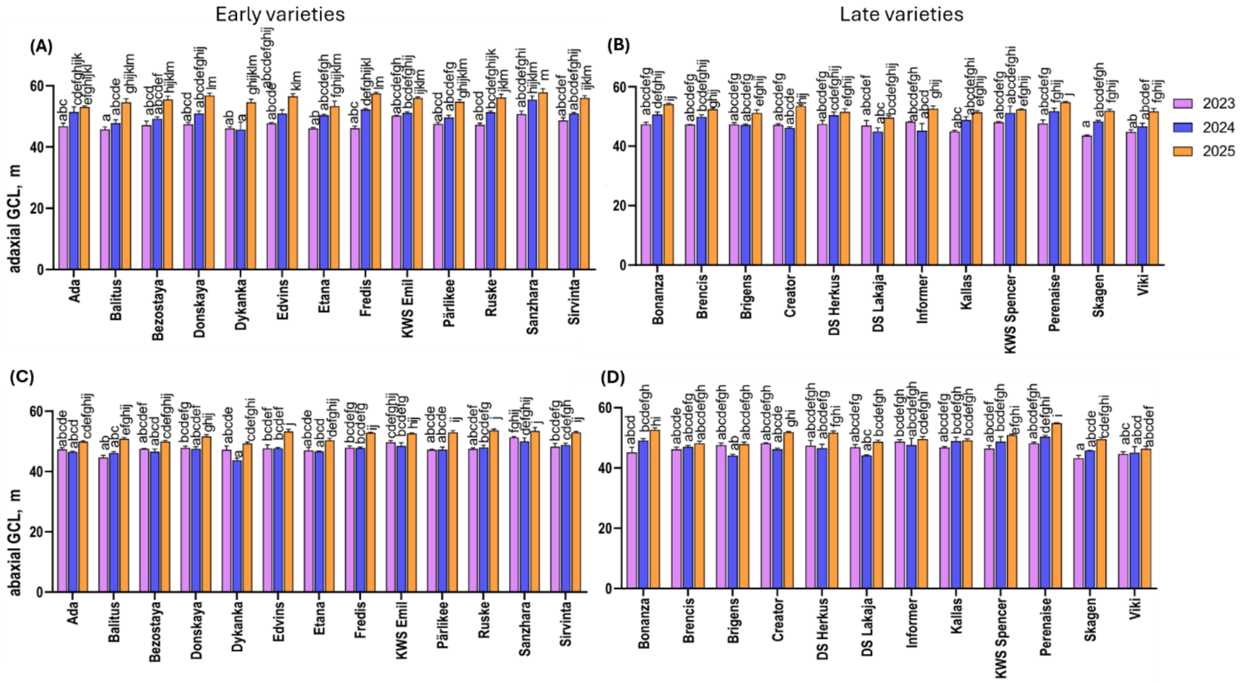


**Figure S7.** Stomatal anatomical traits of winter wheat in trial years. (A) Adaxial guard cell length (GCL, μm) in 2023–2025 of early varieties; (B) Adaxial GCL in 2023–2025 of late varieties; (C) Abaxial GCL in 2023-2025 of early varieties; (D) Abaxial GCL in 2023–2025 of late varieties. Error bars represent standard error of the mean, and different letters indicate statistically significant differences between groups. Tukey's post-hoc test following factorial analysis of variance (two-way ANOVA, p< 0.05, n=3) was used.

Alt text: Graphs A–D. A is showing abaxial guard cell length in early varieties, B is showing abaxial guard cell length in late varieties, C is showing adaxial guard cell length in early varieties, D is showing adaxial guard cell length in late varieties in 2023–2025 with statistical annotations.

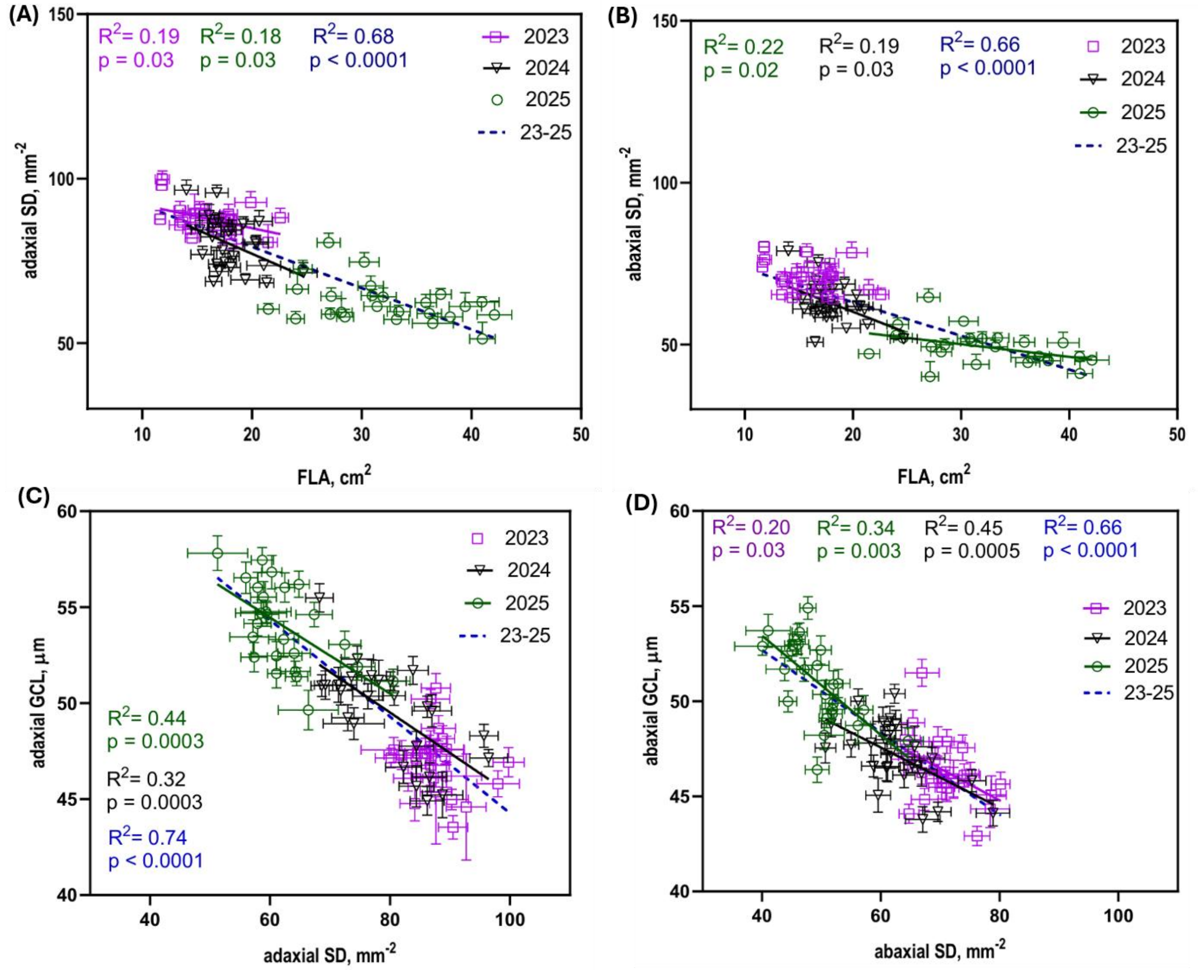


**Figure S8.** Linear regressions between flag leaf area and stomatal traits. (A) Linear regression between flag leaf area (FLA, $cm^2$) and it's adaxial stomatal density (SD, $mm^{-2}$); (B) Linear regression between FLA and it's abaxial stomatal density; (C) Linear regression between abaxial SD and abaxial stomatal GCL; (D) Linear regression between adaxial SD and adaxial GCL. Graphs include linear regression lines for statistically significant relationships ($p < 0.05$, n=25 within each year and n=75 in the pooled dataset, error bars show standard error of the means).

Alt text: Graphs A–D, showing linear regressions between winter wheat stomatal density (SD, $mm^{-2}$), guard cell length (GCL, mm), and flag leaf area (FLA, $cm^2$). All the graphs with significant relationships are with regression lines, statistical significances and p-values.

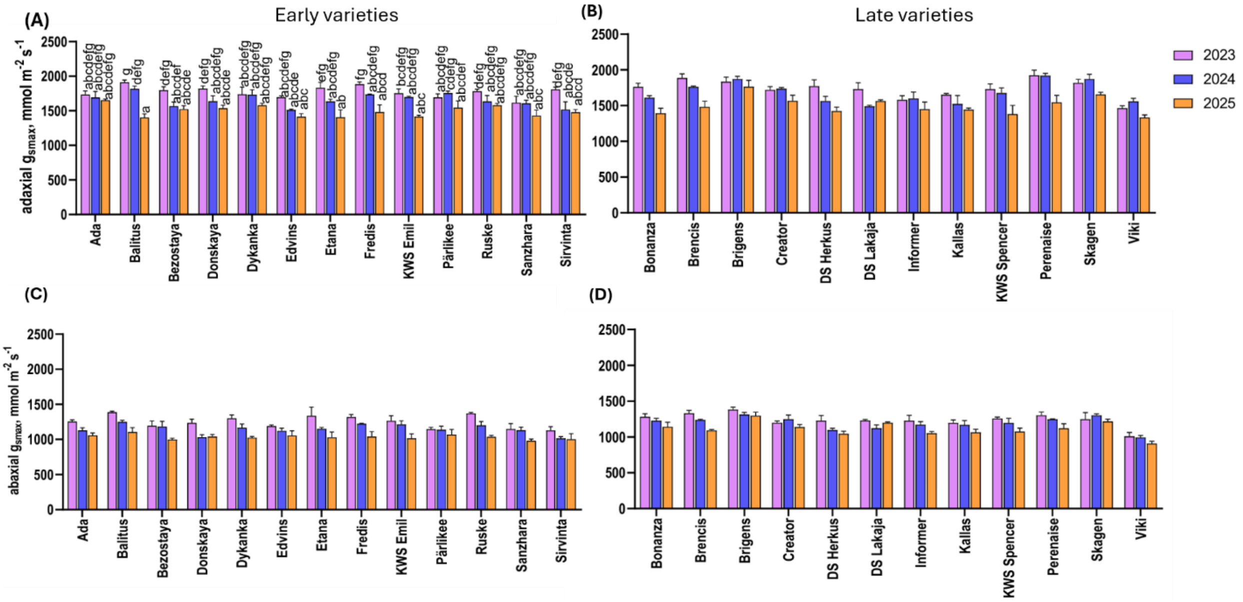


**Figure S9.** Anatomical maximum stomatal conductance ($g_{smax}$, mmol $m^{-2}$ $s^{-1}$) of early and late varieties of winter wheat in 2023–2025. (A) adaxial $g_{smax}$ of early varieties; (B) Adaxial $g_{smax}$ of late varieties; (C) Abaxial $g_{smax}$ of early varieties; (D) Abaxial $g_{smax}$ of late varieties. Error bars represent standard error of the mean, and different letters indicate statistical differences between varieties. Tukey's post-hoc test following factorial analysis of variance (two-way ANOVA, $p < 0.05$, n=3) was used.

Alt text: Graphs A–D, showing winter wheat adaxial and abaxial anatomical maximum stomatal conductance ($g_{smax}$, mmol $m^{-2}$ $s^{-1}$) of early and late varieties in 2023–2025 with statistical annotations.

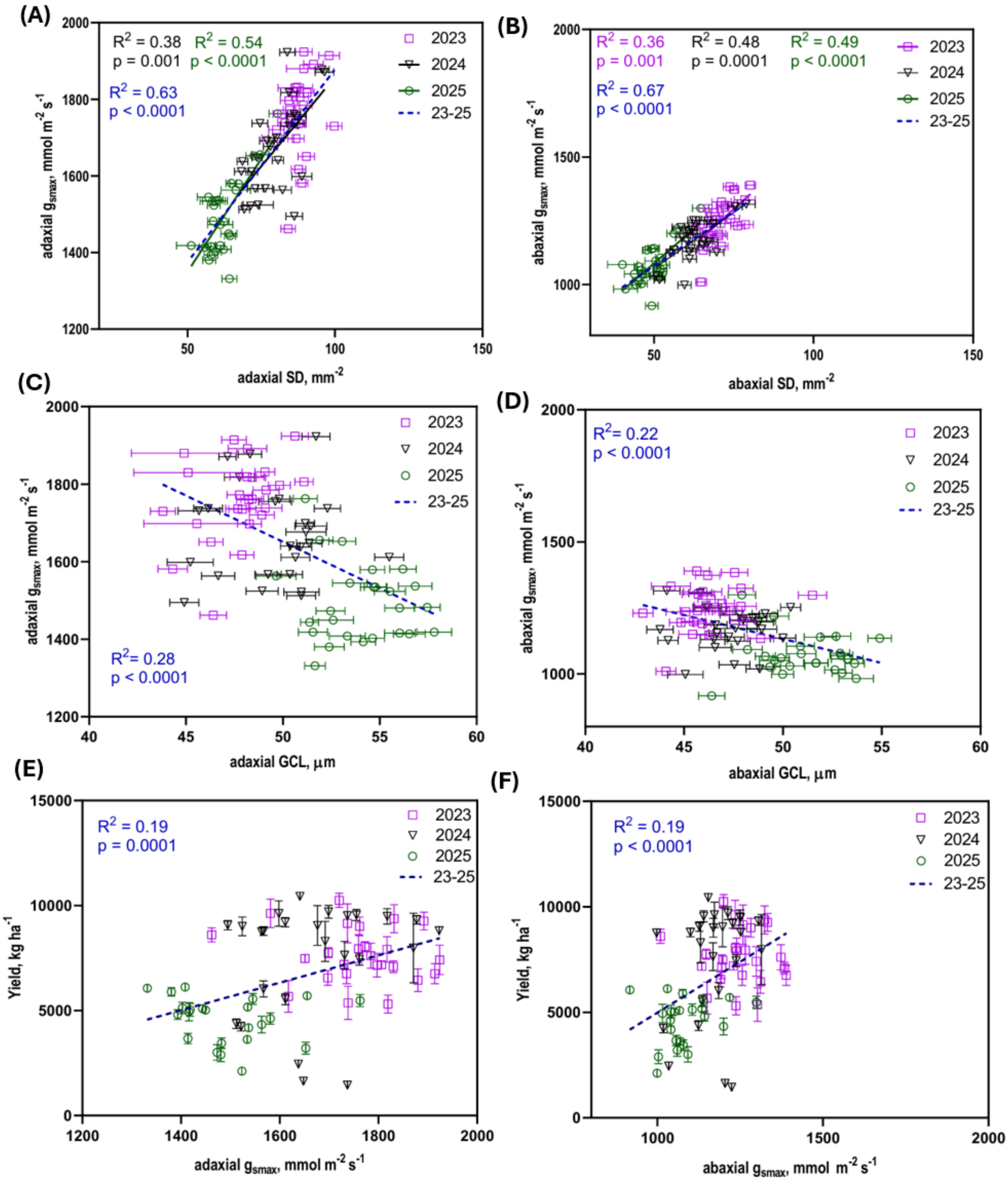


**Figure S10.** Linear regression between stomatal anatomical maximum conductance ($g_{smax}$, mmol m$^{-2}$ s$^{-1}$) and stomatal density (SD, stomata mm$^{-2}$), guard cell length (GCL, μm), and grain yield (kg ha$^{-1}$). (A) Relationship between adaxial SD and adaxial $g_{smax}$ in 2023–2025; (B) Relationship between abaxial SD and abaxial $g_{smax}$ in 2023–2025; (C) Relationship between adaxial $g_{smax}$ and adaxial GCL; (D) Relationship between abaxial $g_{smax}$ and abaxial GCL; (E) Relationship between adaxial $g_{smax}$ and grain yield in 2023–2025; (F) Relationship between abaxial $g_{smax}$ and grain yield in 2023–2025. Graphs with significant relationships include linear regression lines for statistically significant relationships ($p< 0.05$, n=25 for genotypes in each year and n=75 in the pooled dataset, error bars show standard error of the means).

Alt text: Graphs A–F, showing linear regressions between winter wheat stomatal anatomical maximum conductance ($g_{smax}$, mmol m$^{-2}$ s$^{-1}$) and stomatal density (SD, stomata mm$^{-2}$), guard cell length (GCL, μm), and grain yield (kg $ha^{-1}$) in 2023−2025. All the graphs with significant relationships are with regression lines, statistical significances and p-values.

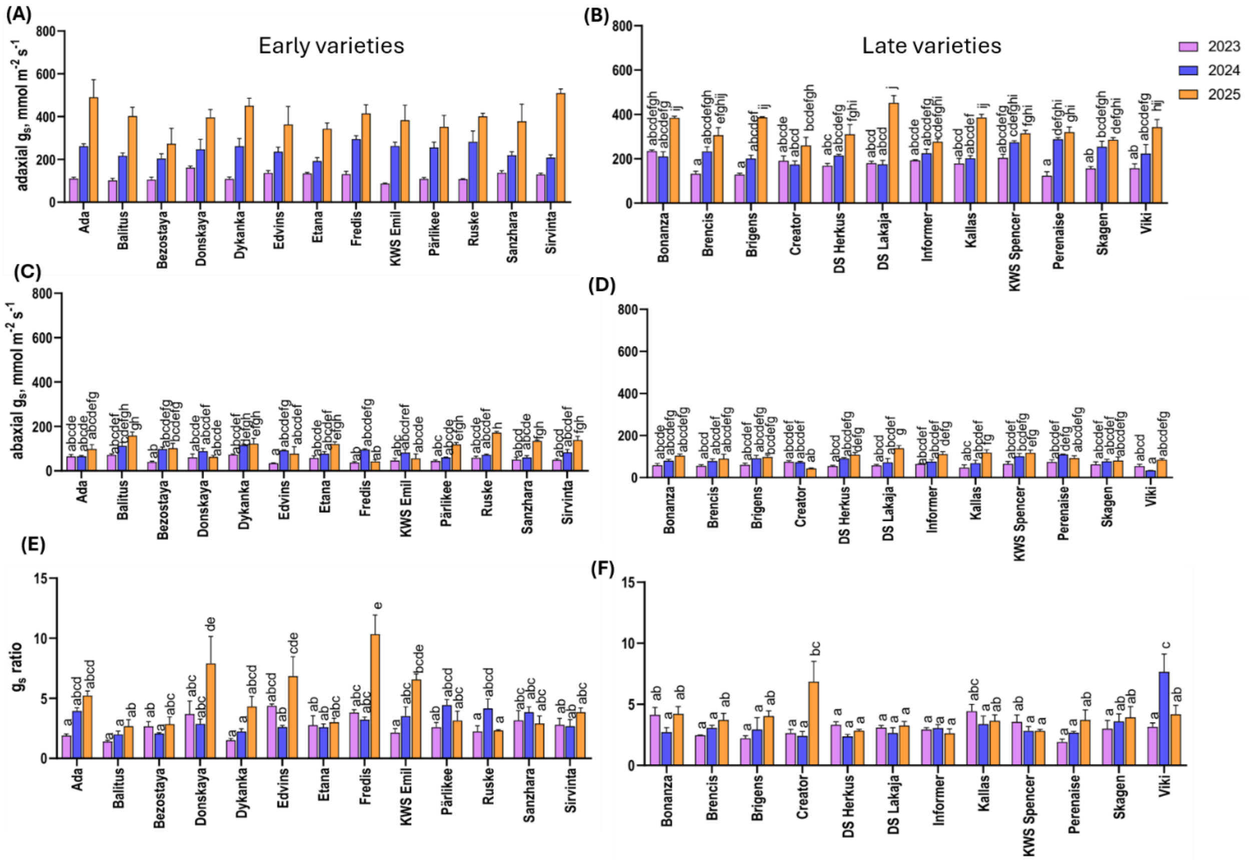


**Figure S11.** Stomatal conductance ($g_s$, mmol $s^{-1}$ $mm^{-2}$) in early and late varieties of winter wheat in 2023–2025. (A) Adaxial $g_s$ in early varieties in 2023–2025; (B) Adaxial $g_s$ in late varieties in 2023–2025; (C) Abaxial $g_s$ in early varieties in 2023–2025; (D) Abaxial $g_s$ in late varieties in 2023–2025; (E) Adaxial to abaxial stomatal conductance ratio ($g_s$ ratio, no unit) in early varieties in 2023–2025; (F) $g_s$ ratio in late varieties in 2023–2025. Error bars represent standard error of the mean, and different letters indicate statistically significant differences between groups. Tukey's post-hoc test following factorial analysis of variance (two-way ANOVA, $p < 0.05$, n=3) was used.

Alt text: Graphs A–F, showing winter wheat stomatal conductance and stomatal conductance ratio in early and late varieties in 2023–2025 with statistical annotations.

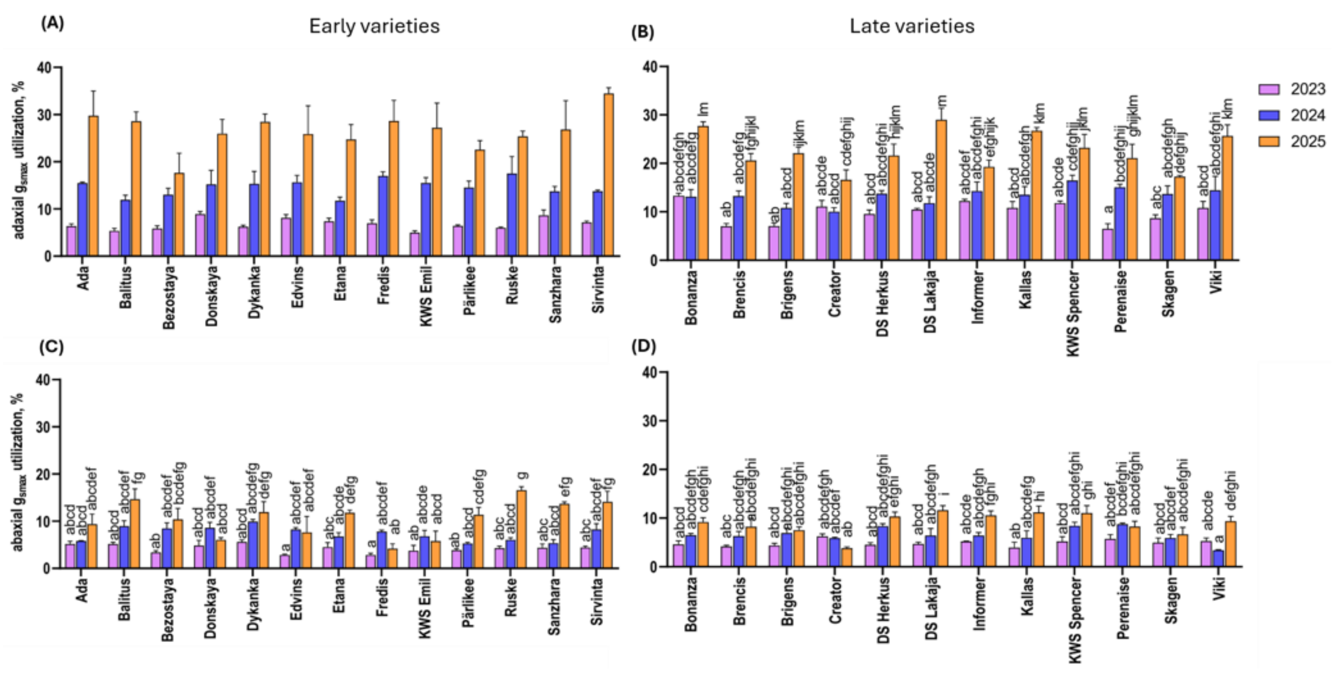


**Figure S12.** Utilization of anatomical maximum stomatal conductance ($g_s$ / $g_{smax}$ *100, %) in early and late varieties of winter wheat in 2023–2025. (A) Utilization of anatomical maximum stomatal conductance on the adaxial side in early varieties ; (B) Utilization of anatomical maximum stomatal conductance on the adaxial side in late varieties ; (C) Utilization of anatomical maximum stomatal conductance on the abaxial side in early varieties ; (D) Utilization of anatomical maximum stomatal conductance on the abaxial side in late varieties. Error bars represent standard error of the mean, and different letters indicate statistically significant differences between groups. Tukey's post-hoc test following factorial analysis of variance (two-way ANOVA, $p < 0.05$, n=3) was used.

Alt text: Graphs A–D, showing Utilization of anatomical maximum stomatal conductance ($g_s$ / $g_{smax}$ *100, %) in early and late varieties of winter wheat in 2023–2025 with statistical annotations.

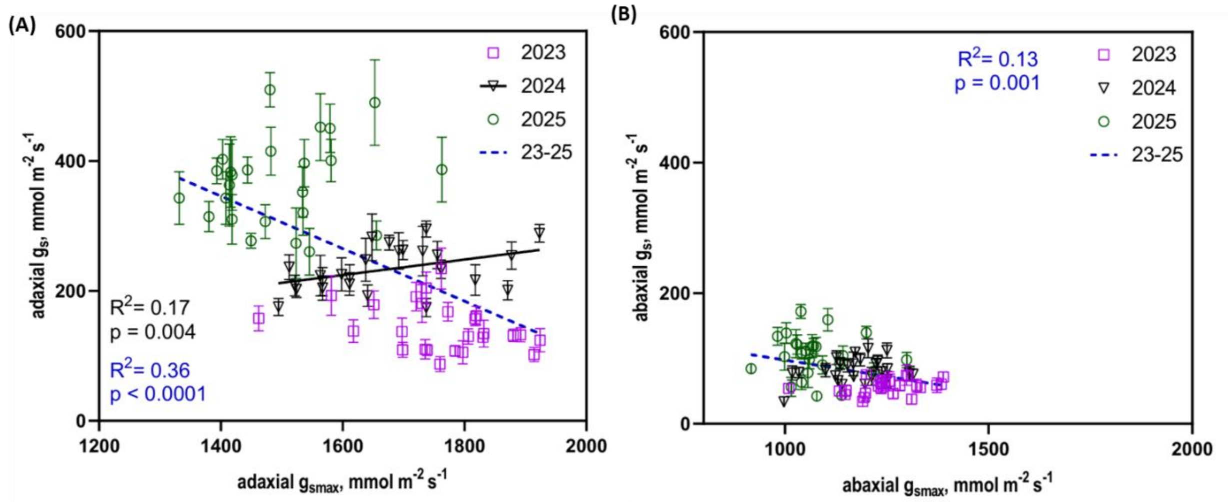


**Fig. S13.** Linear regression between stomatal anatomical maximum conductance ($g_{smax}$, mmol $m^{-2}$ $s^{-1}$) and operational stomatal conductance ($g_s$, mmol $m^{-2}$ $s^{-1}$). (A) Relationship between adaxial $g_s$ and adaxial $g_{smax}$ in 2023–2025; (B) Relationship between abaxial $g_s$ and abaxial $g_{smax}$ in 2023–2025. Graphs with significant relationships include linear regression lines for statistically significant relationships ($p< 0.05$, n=25 for genotypes in each year and n=75 in the pooled dataset, error bars show standard error of the means).

Alt text: Graphs A–B, showing linear regressions between adaxial and abaxial stomatal anatomical maximum conductance ($g_{smax}$, mmol $m^{-2}$ $s^{-1}$) and operational stomatal conductance ($g_s$, mmol $m^{-2}$ $s^{-1}$) of winter wheat in 2023–2025. All the graphs with significant relationships are with regression lines, statistical significances and p-values.
